# Identification of a key genetic factor governing arabinan utilization in the gut microbiome suggests a novel therapeutic target for constipation

**DOI:** 10.1101/2022.12.15.518621

**Authors:** Chengcheng Zhang, Leilei Yu, Chenchen Ma, Shuaiming Jiang, Shunhe Wang, Fengwei Tian, Yuzheng Xue, Jianxin Zhao, Hao Zhang, Liming Liu, Wei Chen, Shi Huang, Jiachao Zhang, Qixiao Zhai

## Abstract

Probiotics have been widely used to improve impaired gastro-intestinal motility, yet their efficacy varied substantially across strains. Here, by a large-scale genetic screen plus *in vivo* measurements, we identified a key genetic factor (*abfA* cluster governing arabinan utilization) in probiotic *Bifidobacterium longum* harnessing the treatment efficacy against functional constipation (FC). Intriguingly, it also presents in a range of gut resident microbiota and played a protective role against FC. Next, our longitudinal multi-omics study in humans revealed that the exogenous *abfA*-cluster- carrying *B. longum* can well establish itself in the gut, and enrich arabinan-utilization residents and beneficial metabolites (e.g., acetate, butyrate, chenodeoxycholic acid and uracil). Finally, transplantation of *abfA*-cluster-enriched human microbiota to FC- induced germ-free mice recapitulated the marked gut-motility improvement and elevated production of beneficial metabolites. Collectively, our proof-of-concept study actively demonstrated a critical yet underexplored role of microbial *abfA* cluster in ameliorating FC, establishing generalizable principles for developing functional-genomics-directed probiotic therapies.

## INTRODUCTION

Functional constipation (FC) is a globally prevalent bowel disorder in humans, with a worldwide prevalence of 10.1% to 15.3% and predominantly seen in women. It’s characterized by recurrent infrequent stools, difficult stool passage, or incomplete defecation ^1, 2^, and associated with cardiovascular disease ^3^, Parkinson disease ^4^, mental disorder ^5^, infantile autism ^6^ and colorectal cancer ^7^. FC imposes a large economic and social burden on patients, impairing their quality of life as well as their ability to work and function socially ^8, 9^.

Impaired gastrointestinal (GI) motility has been implicated with gut microbial dysbiosis, featured by a significant decrease in the abundance of beneficial microorganism (e.g., *Bifidobacteria*, *Lacticaseibacillus*, *Lactiplantibacillus,* etc.) conventionally known as probiotics ^10, 11^. Therefore, the orally-administrated probiotics have been widely used to alleviate FC. Our previous meta-analysis of 15 randomized controlled clinical trials (recruited 1375 participants) reported that probiotic use increased stool frequency by 0.98 bowel movements/week (95% CI: [0.36, 1.60] bowel movements/week) and reduced the whole gut transit time by 13.75 hrs (95% CI: [5.56, 21.93] hrs) ^12^. The introduction of probiotic can lead to changes in end products of gut microbial fermentation such as short-chain fatty acids (SCFAs) which interact with the host immune system and enteric nervous system ameliorating the GI motility ^13^.

However, the therapeutic effect of probiotics against FC often varies substantially among different strains within the same species ^8^. Due to the elusive mechanisms underlying the probiotic strain-specificity, the rational choice of probiotic remains challenging for both medical care professionals and patients ^12, 14, 15^. For FC particularly, the discrepancy in the probiotic efficacy can be mainly explained by two key factors. *i*) What ecological niche a probiotic can occupy in the gut. The stable and robust colonization of the gut of hosts with diverse backgrounds is a prerequisite for probiotic effectiveness. Prior to probiotic ingestion, resident gut microbes have long established their ecological niches which are supposed to exclude most exogenous microbes for a later colonization ^16^. It was documented that unique metabolic capabilities (such as, utilizing dietary fibers poorly utilized by normal residents) usually facilitated the microbial adaptation and stabilized the colonization of a probiotic strain ^16^. *ii*) What functions they can perform. The effective biotherapy using a probiotic strain was supposed to functionally modulate host gut microbiomes (e.g., enhancing the microbial production of SCFAs) or host immunity resulting in significant changes in host physiology ^13^. However, these *in vivo* functional traits can be highly variable across taxonomically identical probiotic strains, and poorly dissected with model organisms from the perspective of both microbial ecology and genomics. Therefore, it is of both scientific and clinical significance to first identify, if any, key genetic elements specific to those functional probiotic strains ^17–20^. Yet, the amount of experimentation necessary to do so is daunting. *i*) To reproduce the strain-specificity issue in the animal model, it is necessary for a research lab to establish a sufficiently large probiotic strain library with paired genomic data, which can cover a high level of genetic variability and link it to their functional differences in *in vivo* gut ecosystem. Numerous studies explored the genomic diversity within a set of probiotic strains, yet, few of them causally linked those strain-specific functional genes to *in vivo* treatment efficacy ^21^ with paired multi-omics study. *ii*) It’s more critical yet challenging to establish such genotype-to-phenotype links in both animal and human models. Most evidence on the beneficial effects of probiotics on gut motility mainly emerged from studies using a mouse model. Probiotic strains were often reported to be effective for a particular disease in animal models yet failed in human clinical trials ^22–24^, or were poorly validated in humans ^25–27^. Therefore, proof-of-concept studies based on the human cohort in combination with evidence from *in vitro* and animal studies is urgently needed for translational research.

This study aimed to identify and characterize the key genetic factor(s) of exogenous probiotics or resident gut microbiota affecting GI motility. We selected the gut native *B. longum* as a model species to identify such genetic factors due to following considerations. *i*) It is the most abundant and one of core members of the human gut *Bifidobacteria* group throughout the human lifespan, showing a strong capability in colonizing the human gut. *ii*) A few *B. longum* strains has been implicated in improving the GI motility in animal models ^28–30^. In this study, *i*) we first established a large library of *B. longum* strains isolated from stool samples of a wide range of Chinese populations. *ii*) The pan-genome analysis of 185 *B. longum* strains was conducted to identify genetically distinct strains, and recapitulated the strain-level functional difference in the probiotic efficacy against loperamide-induced FC in conventionally raised mice. *iii*) A key genetic factor (*abfA* gene cluster preferentially enhancing the arabinan utilization of gut microbes) in the *B. longum* genomes was identified and characterized. *iv*) We next tested the prevalence and functional roles of this genetic factor related to FC in more gut microbial residents (*B. breve, P. pentosaceus*, and *B. nordii*) via the mouse model and at the whole microbial community level by a global, cross-cohort meta-analysis of fecal metagenomics data. To provide the mechanistic insights into how this gut microbial *abfA* cluster works for ameliorating FC, we next conducted a longitudinal multi-omics study (i.e., metagenomics and metabolomics) of fecal samples from human elderly individuals with FC in a clinical trial. The causal relationship between *abfA* cluster and GI motility was further established using a human-to-mouse fecal microbiota transplantation (FMT) experiment, where both *abfA-*cluster-enriched and *abfA-* cluster-depleted human microbiota were transplanted to germ-free (GF) mice with FC (**Figure 1A**). Collectively, we underscored the importance and necessity of characterization of key genetic factors determining the variability in treatment efficacy of probiotics on a particular GI disease.

**Figure 1.**
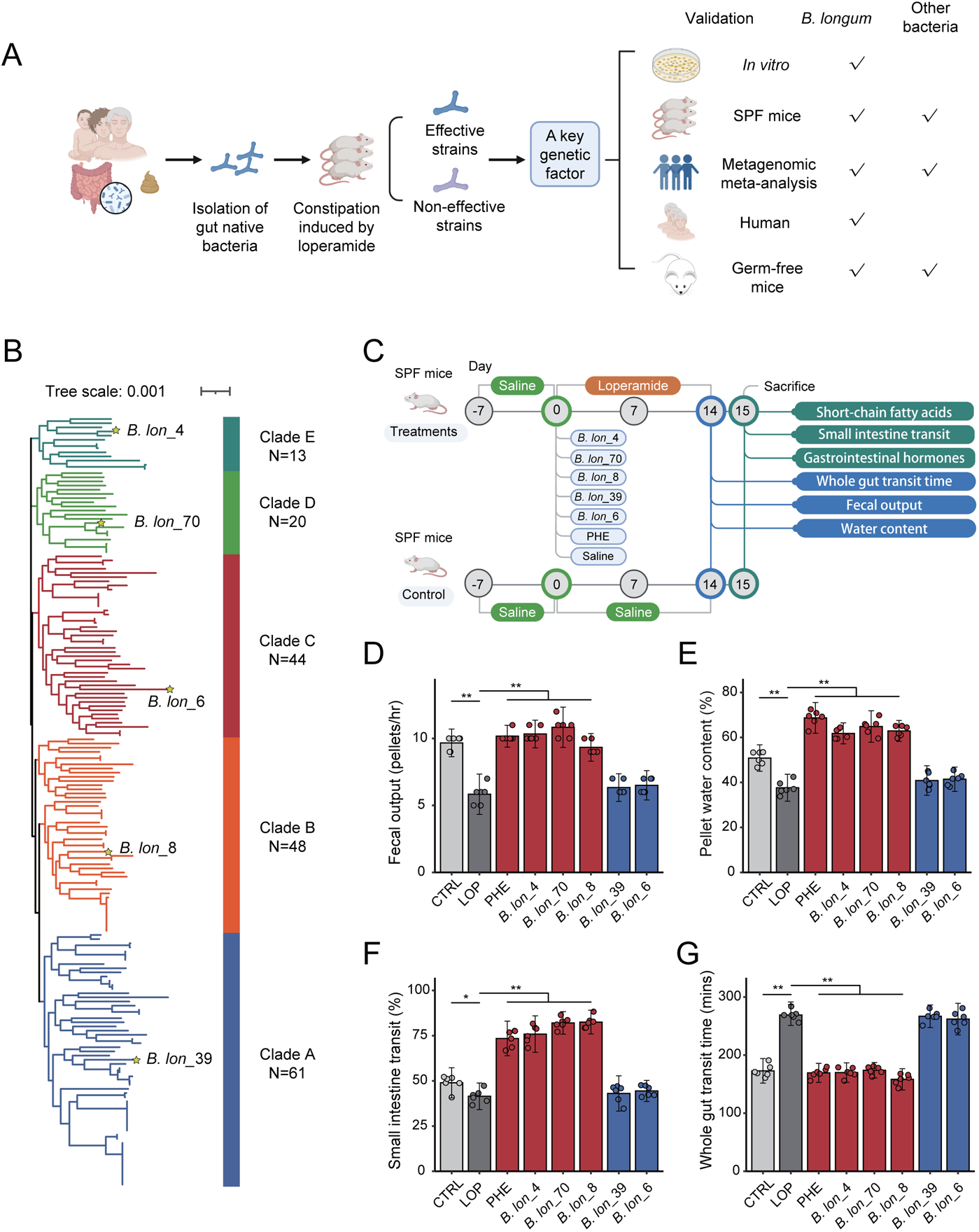
*B. longum* exhibited marked strain-specific effects on constipation induced by loperamide in mice. (**A**) Experimental design for functional-genomics-directed selection strategy of probiotics against FC. “√” represents completion of the test project. (**B**) Whole-genome phylogenetic tree of a representative subset of the five *B. longum* clades comprising the *B. longum* complex. Phylogenetic tree of 185 strains of *B. longum* isolated from our previous research. The tree was built using the maximum likelihood method based on the 821 core genes sequence of *B. longum* strains but visualized by removing the branch length information. The colorful branches indicated the different categorized sub-types in the present research. Five clade-representative *B. longum* strains (*B. lon_*4, *B. lon_*70, *B. lon_*6, *B. lon_*8 and *B. lon_*39) were randomly selected for further animal experiments. (**C**) The experimental design for evaluating the performance of the five randomly selected *B. longum* strains above in alleviating constipation induced by loperamide in SPF BALB/c mice. PHE, phenolphthalein. (**D-G**) The effect of *B. longum* strains on constipation index induced by loperamide in SPF BALB/c mice. Fecal output (**D**), pellet water content (**E**), small intestine transit (**F**), and whole gut transit time (**G**). CTRL, control. LOP, loperamide. N=6 biologically independent animals per group. *p<0.05, **p<0.01 as determined by one-way ANOVA (**D-G**). Data is presented with mean±SD.

## RESULTS

### *B. longum* exhibited strain-specific effect on FC induced by loperamide in mice

To identify genetic factors influencing the probiotic-strain-specific treatment efficacy on FC, we first sought to establish a comprehensive probiotic library that can cover a large genetic and functional diversity of probiotic strains, which enable us to reproduce the widely reported strain-specific effect within our research lab. *B. longum* is known as a major species of native gut bacteria are already adapted to the luminal environment of human hosts, thereby expected to be stably engrafted the human gut in the downstream applications. As a result, a total of 185 new *B. longum* strains were newly isolated from Chinese subjects (N=354) residing in 17 provinces or municipalities in China, aged from 0 to 108 years (**Table S1**). All these new isolates were next whole-genome sequenced for genome analysis for identification of strain-dependent genetic factors. The predicted genome size of these isolates ranged between 2.10 Mb and 2.83 Mb, possessing an average G + C% content of 60.1%, and the number of predicted tRNA genes ranged between 48 to 82 in the majority of strains. We also observed a high genome-wide similarity of 185 *B. longum* strains, namely their average nucleotide identity (ANI) to a reference genome (*B. lon_*8 strain) ranged between 98.02% and 100% (**Figure S2A**). An average of 1949 coding sequences (CDS) per genome was next predicted. The identification of orthologous genes according to the COG (Clusters of Orthologous Groups) database showed that the majority of functional genes in the genome of *B. longum* were involved in housekeeping functions especially those for carbohydrate transport and metabolism (11.34%) as well as amino acid transport and metabolism (10.67%). We identified a pool of 10084 gene families, out of which 821 were highly conservative across these 185 *B. longum* genomes via Roary (**Table S1**). A phylogenetic analysis based on these 821 core genes using the maximum likelihood method clearly divided these 185 *B. longum* strains into five distinct clades (**Figure 1B** and **Table S1**). Such microbial genetic differences often reveal certain strain-specific features and result in diverse phenotypes, which may be associated with host health status ^31, 32^. We reasoned that such a high level of within-strain genetic diversity may lead to the observable divergence in *in vivo* probiotic effect of *B. longum* strains on FC.

Next, we sought to test whether the representative *B. longum* strains (*B. lon_*4, *B. lon_*70, *B. lon_*8, *B. lon_*39 and *B. lon_*6) from the five clades exhibited any divergence in the functionalities, that is, strain-specific effect on treating FC induced by loperamide in a specific pathogen free (SPF) BALB/c mice model (**Figure 1B** and **Figure 1C**). Compared to the control group, mice receiving loperamide treatment showed severe FC symptoms including the reduction in both GI motility and fecal water content (**Figures 1D-1G**). Oral administration of *B. lon_*4, *B. lon_*70 and *B. lon_*8 increased stool frequency, gastrointestinal transit rate and fecal water content, and reduced whole gut transit time (p<0.01, One-way ANOVA test). By contrast, *B. lon_*39 and *B. lon_*6 failed to contain loperamide-induced constipation symptoms in mice under the same conditions (p>0.05, One-way ANOVA test) (**Figures 1D-1G**). We further found that the concentration of fecal acetic acid was remarkably higher in mice engrafted with the effective strains than in those engrafted with the non-effective strains, but not butyric acid, isobutyric acid, pentanoic acid or propionic acid (**Figure S1A**). Notably, effective strains also prevented mice from the loperamide-induced dysbiosis of gastrointestinal hormones (VIP, Gas and MIL) in the blood stream (**Figure S1B**). Collectively, we reproduced the probiotic strain-specificity in the treatment efficacy against FC in SPF BALB/c mice using the clade-representative *B. longum* strains from a large pool of candidate probiotics (N=185). Furthermore, the engraftment of effective probiotic *B. longum* drastically attenuated FC through elevated production of acetate in the gut and modulating blood-derived hormones.

### The effective *B. longum* strains harbored two specific gene clusters (*abfA* and *hypBA*) both encoding arabinofuranosidases

To gain genetic insight between effective strains (*B. lon_*70, *B. lon_*4 and *B. lon_*8) and non-effective strains (*B. lon_*6 and *B. lon_*39), we performed comparative genomic analysis of these five representative *B. longum* strains. The ANI among these five complete *B. longum* genomes ranged from 98.4% to 99.1% (**Figure S2A**), indicating that functional differences (e.g., differences in the probiotic treatment efficacy) between strains is not necessarily related to the global pattern of nucleotides in the genomes but probably due to as low as 1% genetic difference. Using a minimum sequence identity threshold of 95% for orthologous genes, we identified five syntenic loci (or gene clusters) that are present in the effective *B. longum* strains but absent from the non-effective ones (**Table S2**). An *adfA* gene cluster (or *abfA* cluster below) contained four linked genes that were all functionally annotated as α-L-arabinofuranosidase (*abfA*) at loci 1 were specific to the effective *B. longum* strains. At loci 4, we also found an effective*-B. longum*-specific *hypBA* gene cluster (*hypBA* cluster) including six genes encoding β-galactosidase, β-L-arabinobiosidase, β-L-arabinofuranosidase and three ABC transporter permease (**Figure 2A and Figure 2B**). At the other three loci (loci 2, 3, and 5), by contrast, paralogous genes were also present in the genomes of the non-effective *B. longum*, which were thus not included in this study (**Table S2**). These two gene clusters are functionally related to degrading dietary arabinan, a significant portion of pectic polysaccharides in various plant tissues ^33, 34^.

**Figure 2.**
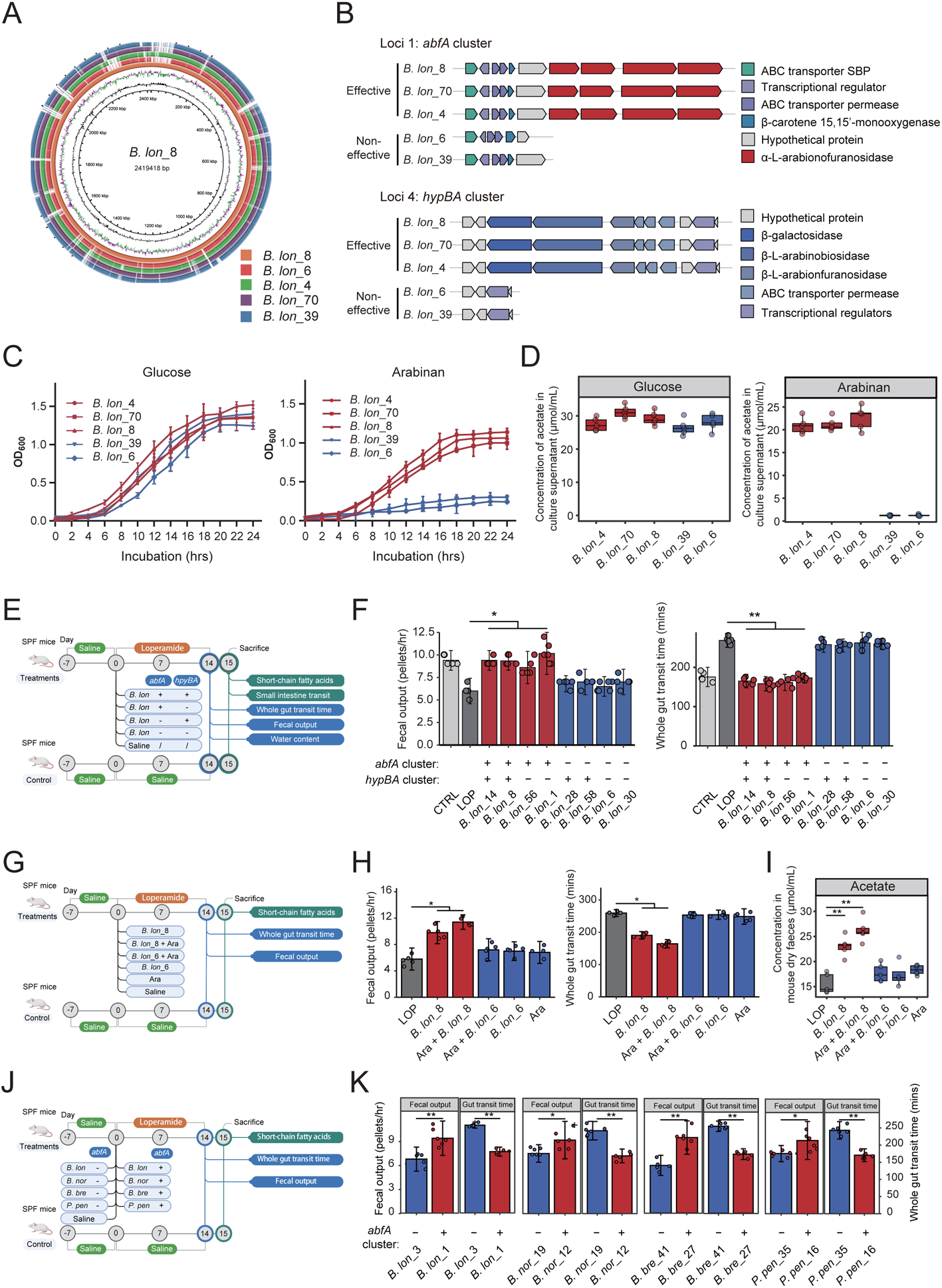
The *abfA* cluster is a keystone functional element for gut microbiome-mediated alleviation of constipation. (**A**) Genomic comparison of the chromosomes of five strains. Both sequences are started from the predicted replication origin. From inner to outer: (1) GC Content, (2) GC Skew, (3) *B. lon_*8, *B. lon_*6, *B. lon_*4, *B. lon_*70 and *B. lon_*39. (**B**) Genomic regions containing genes encoding specific arabionofuranosidases in the sequenced *B. longum* strains. Genes and their orientations are depicted with arrows. (**C**) Growth curve of *B. longum* strains (*B. lon_*8, *B. lon_*6, *B. lon_*4, *B. lon_*70 and *B. lon_*39) in media containing glucose or arabinan. The strain was cultured at 37°C in sugar-restricted basal medium supplemented with 2.0% different carbon sources under anaerobic conditions. The OD_600_ was measured every 2 hrs for 24 hrs. Error bars indicate SD (N=3). (**D**) Concentrations of acetate in culture supernatant after 24-hr incubation with addition of glucose or arabinan as a substrate, as determined by GC/MS. (**E**) The experimental design for evaluating the probiotic performance of *B. longum* strains that harbors *abfA* gene cluster, *hypBA* gene cluster, or both. The mice were treated with *B. longum* strains (1×10^9^ CFU per day) for 14 continuous days. “+” denotes that the gene cluster is present. “-” denotes that the gene cluster is absent. (**F**) Constipation related index from SPF BALB/c mice treated with *B. longum* strains harbor *abfA* gene cluster, *hypBA* gene cluster, or both. (**G**) The experimental design for evaluating the probiotic performance of the *B. longum,* or arabinan, or *B. longum* plus arabinan for 14 continuous days. Ara, arabinan. (**H**) Constipation index from mice after administration of *B. longum* strains, with or without arabinan, or only arabinan. (**I**) Concentrations of acetate in the mouse feces at day 14 after *B. longum* strain or/and arabinan treatment, as determined by GC/MS. (**J**) The experimental design for evaluating the probiotic performance of *B. longum*, *B. breve*, *P. pentosaceus* and *B. nordii* strains with or without the *abfA* gene cluster. (**K**) Effect of bacteria strains on constipation index induced by loperamide in mice. *p<0.05, **p<0.01 as determined by one-way ANOVA (**F, H and I**), Wilcoxon rank-sum test or Student’s *t* test (**K**). N=5-6 biologically independent animals per group (**F-K**).

We next returned these effective and non-effective strains back to *in vitro* condition, and tested if such strain-level genetic differences directly resulted in distinct arabinan-utilization capability. Interestingly, in our *in vitro* experiment, the effective strains (i.e., *B. lon_*4, *B. lon_*70 and *B. lon_*8) grew substantially faster than non-effective strains (*B. lon_*39 and *B. lon_*6) in media containing arabinan in a 24-hr incubation (**Figure 2C**). Furthermore, effective strains drastically enriched the production of acetic acid (>20 μmol/mL), the key end product of probiotic *Bifidobacterium* in the carbohydrate synthesis and metabolism, by the efficient utilization of arabinan, while non-effective strains didn’t produce acetic acid at a meaningful level (<1 μmol/mL, **Figure 2D**). By contrast, we didn’t observe any difference in the growth (**Figure 2C**) or production of acetic acid (**Figure 2D**) between effective and non-effective strains in media containing glucose along the whole 24-hr incubation. Next, we examined the *in vitro* transcription-level activity of each gene in these two gene clusters specific to effective strains under either arabinan or glucose incubation. It was reported that arabinan can increase the expression level of gene *BLLJ_1852* and *BLLJ_1853* in a similar gene cluster (*abfA*) of *B. lon*gum JCM1217 ^33^. In our study, arabinan induced a more extensive gene transcription profile in *abfA* cluster than reported. It upregulated the expression of eight out of ten genes (*BLLJ_1850*, *BLLJ_1852*, *BLLJ_1853*, *BLLJ_1855*, *BLLJ_1856*, *BLLJ_1857*, *BLLJ_1858* and *BLLJ_1859*) in the *abfA* cluster of *B. lon_*8, whereas only three out of 11 genes in the *hpyBA* cluster showed a slightly increased expression level (**Figure S2B and Figure S2C**). In the *abfA* cluster, the relative mRNA expression to 16S rRNA gene (%) was averagely 0.35, while it was only 0.019 averagely in the *hpyBA* cluster. In terms of the level and breath of gene expression, *abfA* cluster made an overwhelmingly greater contribution to arabinan utilization than *hpyBA* cluster in the presence of arabinan. From the evidence from *in vitro* bacterial growth experiment, metabolomic analysis, genomic analysis, and mRNA expression profiling, the probiotic properties of effective *B. longum* strains would largely attribute to the efficient utilization of the plant-derived pectic polysaccharides by the active involvement of *abfA* cluster.

### The *abfA* cluster is solely responsible for gut microbiome*-*mediated alleviation of FC

We next sought to test whether *abfA* cluster or *hypBA* cluster in the *B. longum* genome can independently influence the *in vivo* treatment efficacy on FC. It’s well known that *B. longum* is reluctant to be genetically manipulated (e.g., knocking out a gene cluster) for generating a mutant library. Given the diverse genotypes of *B. longum* presented in our large strain library, we reason that we should employ the naturally occurring mutants to separate the genetic contribution of *abfA* and *hypBA* clusters to the functional traits in the *in vitro* and animal models. Tracing back to original 185 *B. longum* strains in the library, 105 (56.8%) possessed a complete *abfA* cluster, while 129 (69.7%) carried a complete *hypBA* cluster. Interestingly, 79 out of 185 strains (42.7%) possessed both gene clusters (**Figure S2D**). From this strain library, eight *B. longum* strains with distinct presence/absence patterns of *abfA* cluster and *hypBA* cluster in their genomes were selected to conduct the animal experiments. These strains can be classified into four groups: *abfA+hypBA*+ (i.e., *B. lon_*14, and *B. lon_*8), *abfA+hypBA*- (i.e., *lon_*56 and *B. lon_*1), *abfA-hypBA*+ (i.e., *B. lon_*28 and *B. lon_*58), *abfA-hypBA*- (i.e., *B. lon_*6 and *B. lon_*30) (**Figure 2E)**. *In vitro* growth test of arabinan as the only carbon source showed that strains carrying an *abfA* cluster grew exponentially faster than others after a 6-hr incubation, indicating that *abfA* cluster was likely a key driver for the microbial growth through the active arabinan metabolism in *B. longum* (**Figure S3A and Figure S3B)**. Furthermore, those *B. longum* strains carrying *abfA* cluster (*B. lon_*14, *B. lon_*8, *B. lon_*56 and *B. lon_*1) consistently increased fecal output and decreased whole gut transit time (p<0.05, **Figure 2F**), whereas the strains encoding the *hypBA* cluster alone (*B. lon_*28 and *B. lon_*58) showed no effects (p>0.05, **Figure 2F**) on reversing FC symptoms. Therefore, we reasoned that *abfA* cluster should be a sole genetic factor determining probiotic efficacy of *B. longum* against FC. To validate the above findings, we additionally collected more strains from the library, i.e., *B. lon_*32 (*abfA+hypBA*+), *B. lon_*63 (*abfA+hypBA*-), and *B. lon_*64 (*abfA-hypBA*-) and performed the same *in vitro* experiments and animal experiments. *B. lon_*32 and *B. lon_*63 consistently increased fecal output and decreased whole gut transit time (**Figures S3C-S3F**) over time, whereas *B. lon_*64 showed no significant effects (**Figures S3C-S3F**), further demonstrating the key role of *abfA* cluster in ameliorating FC through the enhanced production of acetate.

Given *abfA* cluster specialized in arabinoglycan utilization, we next investigated the synergistic effect of effective *B. longum* strains and dietary arabinan in mice with FC (**Figure 3G**). Firstly, arabinan treatment alone did not affect the fecal output and whole intestinal transit time (**Figure 3H**). It suggested that native gut microbiota of mice was not able to assimilate arabinan probably due to certain genetic deficiency. Moreover, supplementation of arabinan with the *abfA*-cluster-carrying *B. lon_*8 significantly increased fecal output and reduced whole intestinal transit time than did *B. lon_*8 alone (**Figure 3H**). By contrast, *abfA*-cluster-deficient *B. lon_*6 showed no effects on FC symptoms even when sufficient dietary arabinan was added. The GC-MS analysis of fecal samples further revealed that the supplementation of *B. lon_*8 resulted in an increase in the production of acetate in the gut than *B. lon_*6. Supplementation of both dietary arabinan and *B. lon_*8 significantly promoted the acetate production which has been widely implicated in increasing the GI motility (**Figure 2I**). Overall, the *abfA* cluster was a critical genetic determinant in promoting arabinan assimilation and acetate production in the gut, which effectively increased GI motility in mice.

**Figure 3.**
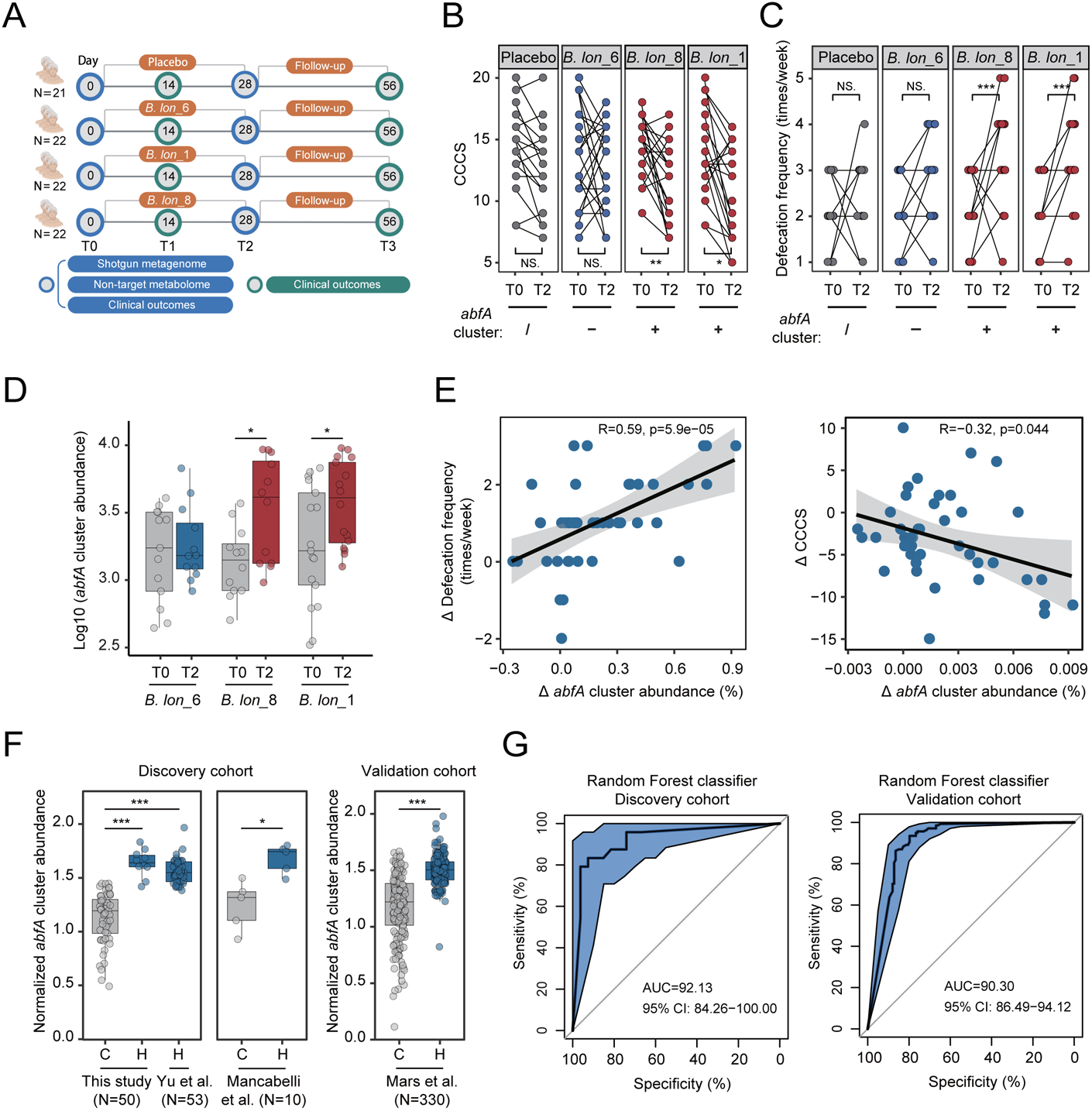
The protective role of *abfA* cluster in the gut microbiota against constipation can be validated in the multiple human cohorts. (**A**) The experimental design. A total of 87 elderly individuals were randomly divided into four groups through Medidata Balance methods: Placebo group, *B. lon_*6 group, *B. lon_*8 group and *B. lon_*1 group. Shotgun metagenome and non-target metabolome were performed to explore the microbiome and metabolites profile of fecal samples, and the clinical indexes were also determined in each time point. Only individuals with paired data were included in each analysis. T0, day 0; T1, day 7; T2, day 28; T3, day 56. (**B**) The differential distributions of CCCS in individuals before and after 28 days of *B. longum* supplementation. (**C**) The differential distributions of defecation frequency in individuals before and after 28 days of *B. longum* supplementation. (**D**) The differential *abfA* gene cluster abundance in individuals after 28 days of *B. longum* strains supplementation. (**E**) The left panel, the correlation between the change in defecation frequency and the change in *abfA* gene cluster abundance from baseline to 28 days after *B. longum* strains supplementation. The right panel, the correlation between the change in CCCS and the change in *abfA* gene cluster abundance from baseline to 28 days after the supplementation of *B. longum* strains. A linear model (LM) was used for creating a smoothed line for scatters. The grey area around the smoothed line indicates the 95% confidence interval. The R and p value in the scatter plots were calculated from Spearman correlation analysis. (**F**) The relative abundances of *abfA* gene cluster across all subjects in our study (N=50), Yu et al. (N=53), Mancabelli (N=10), and Mars et al. (N=330) cohorts. Overall, the relative abundance of *abfA* gene cluster is significantly higher in the health control group compared to the functional constipation patients. (**G**) The receiver operating characteristic (ROC) curve and the area under curve (AUC) exhibited excellent performances both in the training and validation data, which was calculated based on the normalized *abfA* gene cluster abundance. The error bands indicate the 95% confidence intervals. Discovery cohort datasets, N=113; validation cohort datasets, <0.05, **p<0.01, ***p[<0.001 as determined by paired Student’s *t* test (**B-D**) or Wilcoxon signed-rank test (**F**).

Other than *B. longum*, a few common commensal gut microbes also possess a complete *abfA* cluster, including *Bifidobacterium breve* (*B. bre*)*, Pediococcus pentosaceus* (*P. pen*) and *Bacteroides nordii* (*B. nor*). To test whether the *abfA* cluster non-specific to *B. longum* can also deliver its key probiotic function: improving gut motility, we next compared the treatment performance of *B. longum* strains and three representative gut residents either carrying or lacking an *abfA* cluster (**Figure 2J**). Consistent with our observations in *B. longum*-related experiments, all gut commensal organisms carrying the *abfA* cluster (*B. lon_*1, *B. bre*_27, *B. nor_*12 and *P. pen_*16) remarkably increased fecal output and decreased whole gut transit time, whereas the *abfA-*cluster-deficient strains (*B. lon_*3, *B. bre*_41, *B. nor_*19 and *P. pen_*35) did not show any potentials in improving the bowel movement (**Figure 2K**).

We next measured the concentration of three SCFAs (acetate, butyrate and propionate) before and after treatment in each group. Despite the consistent phenotypic changes observed in mice administrated with *abfA-*cluster-carrying *B. longum*, *B. breve*, *P. pen* and *B. nor*, the temporal change in the production of SCFAs was strain-specific. Both effective *B. longum* and *B. breve* significantly increased the concentration of acetic acid in feces. Effective *P. pen* significantly increased the content of butyric and propionic acid, while effective *B. nor* increased the content of acetic and propionic acid in feces (**Figures S3I-S3K**). Intriguingly, we found that the functional role of *abfA* cluster against FC is totally not dependent on specific gut microbes. It would be rather like a universal genetic element in the gut microbiota influencing the host constipation phenotype. To our knowledge, this gene cluster plausibly facilitated these “carrier” commensal microbes (*B. longum*, *B. breve*, *P. pen* and *B. nor*) to overcome the colonization resistance issue through acquiring a unique capability in utilization of arabinan, which enabled them to establish their ecological niches. Then these well gut-adapted microbes can produce beneficial yet specific SCFAs commonly promoting GI motility.

### Validation of the protective roles of *B. longum* equipped with *abfA* cluster against FC in a Chinese elderly cohort

To further validate the protective role of *abfA* cluster against FC in the human cohort, we selected three strains of *B. longum* either possessing an *abfA* cluster (*B. lon_*8 and *B. lon_*1) or possessing no *abfA* cluster (*B. lon_*6) in the genome for FC patients. We conducted a double-blind, randomized, placebo-controlled clinical trial. A total of 120 participants in this study were recruited from the Affiliated Hospital of Jiangnan University, 33 participants excluded. Finally, 87 elderly individuals diagnosed with FC were enrolled in the trial after completing a standardized protocol for initial hospital-based management of FC. These participants were subsequently randomly assigned to one of four treatment groups and underwent randomization to receive once daily a placebo (2 g maltodextrin per day, N=21), *B. lon_*6 (1×10^9^ CFU per day, N=22), *B. lon_*8 (1×10^9^ CFU per day, N=22) or *B. lon_*1 (1×10^9^ CFU per day, N=22) supplementation for 28 days (**Figure 3A**). All 87 subjects in the clinical trial completed the study. Distribution of sex, age, BMI and Cleveland Clinical Constipation Score (CCCS), and defecation frequency, among other parameters, of the study participants were not statistically significant different among the four subject groups at baseline (**Table S3**).

Similar to above observations in animal experiment, *B. lon*_1 and *B. lon*_8 increased stool frequency on average by 1.04 bowel movements/week (95% CI: [0.62, 1.51] bowel movements/week, **Table S3**) and 1.15 bowel movements/week (95% CI: [0.66, 1.64] bowel movements/week, **Table S3**) from baseline to day 28, respectively. *B. lon*_1 and *B. lon*_8 reduced CCCS on average by 4.09 (95% CI: [2.13, 6.05], **Table S3**) and 4.86 (95% CI: [3.00, 6.36], **Table S3**) from baseline to day 28, respectively. In contrast, no significant differences were observed in placebo and *B. lon_*6 group where the *abfA*-cluster-deficient probiotic *B. longum* was administrated (**Figure 3B** and **Figure 3C**). The *B. lon_*1 and *B. lon_*8 supplementation also resulted in a significant reduction in Patient Assessment of Constipation Symptoms (PAC-SYM) (− 0.40±0.56 in *B. lon*_1 group and −0.45±0.40 in *B. lon*_8 group, respectively; **Table S3**) and in Patient Assessment of Constipation Quality of Life global score (PAC-QoL) (− 0.42±0.44 in *B. lon*_1 group, and −0.37±0.27 in *B. lon*_8 group, respectively; **Table S3**) from baseline to the end of intervention (i.e., from baseline to day 28). At the follow-up visit (day 56), we observed a persistent reduction in the FC symptoms (PAC-SYM, PAC-QoL) resulting from the supplementation of the *B. lon_*1 or *B. lon_*8, despite that the probiotics was not administrated from day 28 to 56 (**Table S3**). Moreover, rectal, abdominal, stool and physical discomfort decreased significantly at the end of intervention compared to baseline in *B. lon_*1 and *B. lon_*8 group (see **Table S3** for clinical characteristics). We then examined the impact of effective *B. longum* strains on overall abundance of the microbial *abfA* cluster in feces using shotgun metagenomic sequencing. The administration of *B. lon_*1 and *B. lon_*8 remarkably increased *abfA*-cluster abundance in the gut microbiota at day 28 from baseline (**Figure 3D**, whereas *abfA*-cluster-deficient *B. lon_*6 supplement didn’t (**Figure 3D**). The temporal changes in *abfA* cluster abundance positively correlated with those in defecation frequency (r=0.59, p<0.0001, Spearman’s rank correlation) and negatively correlated with those in CCCS (r=-0.32, p<0.05, Spearman’s rank correlation, **Figure 3E**). However, temporal shifts in the overall abundance of *B. longum* didn’t significantly correlate with those in defecation frequency or CCCS (**Figures S4A-S4D**). It indicated that *abfA* cluster in gut microbiome rather than its carriers (i.e., *B. longum* strains) correlated with host phenotype. If this holds true, then *B. longum* strains weren’t the main source species of *abfA* cluster in the gut microbiome, and moreover, this gene cluster may spread to more resident species afterward. It is plausible that the ingested *abfA*-cluster-carrying *B. longum* strains can elicit more gut residents carrying the *abfA* cluster thriving in the gut microbial community through the unique metabolic capability on the arabinan utilization. Then, we reason that this gene cluster represents a key protective microbiome function against FC and would necessarily be differentially abundant in the gut microbiota between healthy individuals and FC patients, yet largely neglected by previous fecal metagenomics studies.

### The global multi-cohort metagenomic meta-analysis revealed a strong predictive power of *abfA* cluster abundance in human gut microbiome in predicting FC

Here, we additionally analyzed 330 fecal metagenomics samples from Mars et al ^35^ and 10 samples from Mancabelli et al ^36^ to validate and support our hypothesis. To increase our sample size in healthy controls, we also examined 53 metagenomic samples from the Yu et al ^37^. Compared to healthy individuals (N=10, this study; N=53, Yu et al.), the abundance of *abfA* cluster in fecal metagenomics of patients with FC (N=40) was significantly lower (p<0.001, Wilcoxon rank-sum test, **Figure 3F**). Consistently, a lower abundance of *abfA* cluster in FC patients was observed in both Mancabelli et al. and Mars et al. cohorts compared to healthy individuals (p<0.05 and p<0.001, respectively; Wilcoxon rank-sum test, **Figure 3F**). Next, we trained a supervised classification model for distinguishing FC and healthy controls based on the normalized *abfA* cluster abundance using Random Forests. The predictive model exhibited a high classification accuracy (area under the receiver operating characteristic curve, or AUC=92.13%, 95% CI: 84.26−100.00%, N=122, **Figure 3G**) in the discovery cohort (including subjects from this study, Yu et al. and Mancabelli et al. studies), which was well validated in an external cohort (Mars et al, AUC=90.30%, 95% CI: 86.49−94.12% N=330, **Figure 3G**). It suggested that abundance of this single gene cluster estimated from metagenomic data can be possibly developed as a simple and powerful biomarker to either diagnose FC or predict the therapeutic outcomes of FC.

Due to the intrinsic link between *abfA* cluster and high-fiber diet, we next hypothesized a relevance of this functional signature to a broader range of human chronic diseases associated with gut microbial dysbiosis. Next, we estimated the normalized abundance of *abfA* cluster in a panel of 11 metagenomic datasets (N=1724) linking to multiple disease types, including colorectal cancer (CRC, N=269) ^38–41^, chizophrenia (SCZ=171) ^42^, grave’s disease (GD, N=164) ^43^, type 2 diabetes (T2D, N=145) ^44^, breast cancer (BC, N=133) ^45^, inflammatory bowel diseases (IBD, N=220) ^46^, liver cirrhosis (LC, N=237) ^47^, and atherosclerotic cardiovascular disease (ACVD, N=385) ^48^ (**Table 1**). Interestingly, we found that this single gene cluster distinguishes ACVD, T2D and CRC from health controls respectively and presented in a lower abundance in cases that controls in each cohort (Wilcoxon rank-sum test, p<0.001 in ACVD, p<0.05 in T2D and p<0.05 in CRC, **Figure S5**). Since these human diseases have been broadly linked to a lack of high-fiber diet for patients, it suggests that the *abfA* cluster would be a promising gut microbiome signature associated with a range of human diseases result from a limited capability to digest and absorb fiber-derived nutrients, such as arabinans.

**Table 1.**
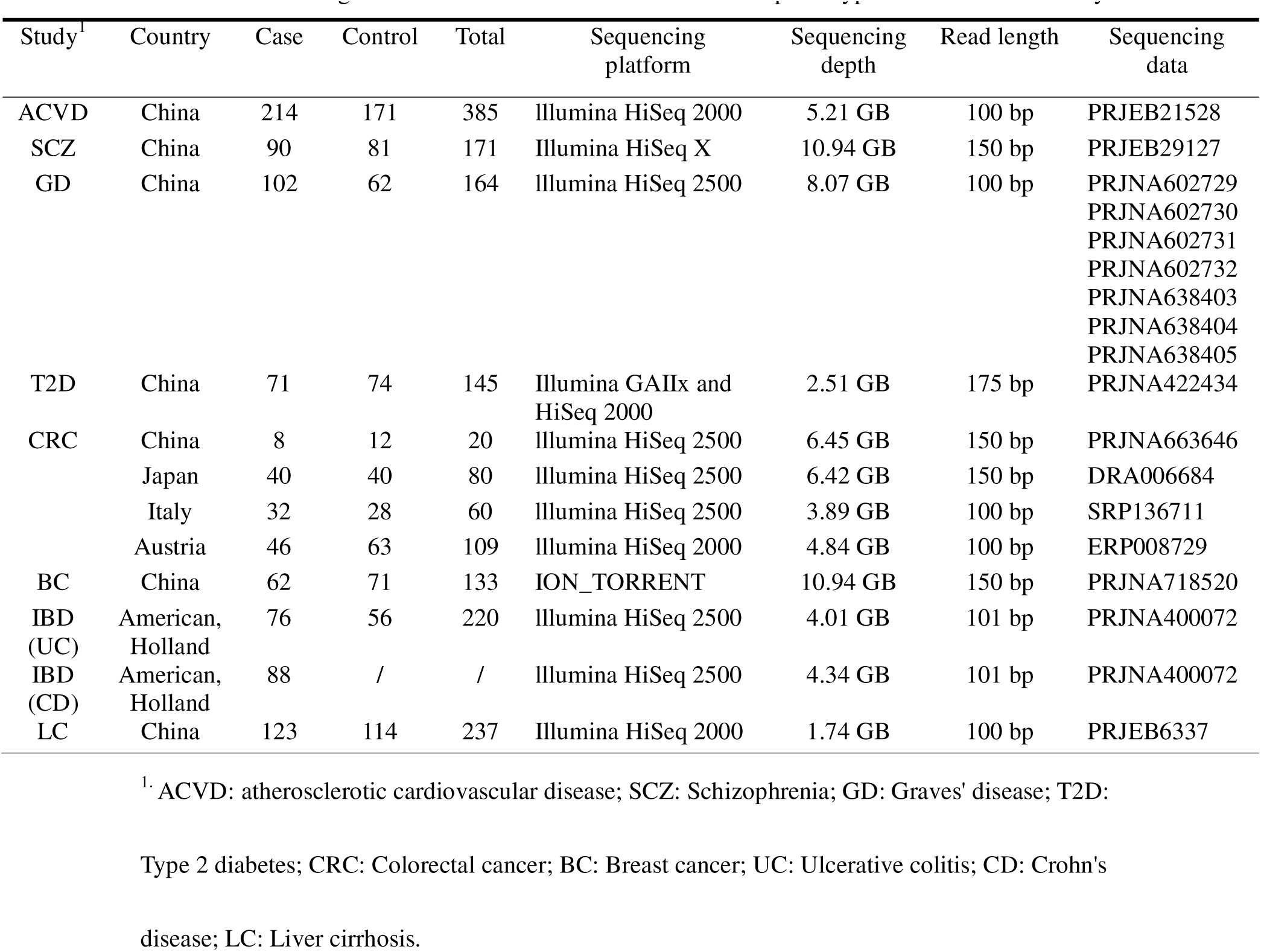
Fecal metagenomic studies of different human disease phenotypes included in this study

### The shifts in specific gut resident strains responsive for the elevated abundance of intestinal microbial *abfA* cluster after *B. longum* administration in humans

To explore the ecological effect of elevating the abundance of microbial *abfA* cluster (**Figure 3D**), we investigated the alterations in diversity and composition of the resident gut microbiota in our cohort. Here, we first tested if the microbiome temporally differed before and after *B. longum* administration for each host using Bray-Curtis dissimilarity. Interestingly, at either genus or species level, no significant within-host microbiome difference in the fecal microbial community of individuals from baseline was observed for any host groups (p>0.05, PERMANOVA, **Figure 4A, Figure S6A and Figure S6B**). We further asked if probiotic-induced ecological changes in the resident microbiota would be discernable at a finer taxonomic resolution, such as the strain level. We then assembled all the shotgun metagenomic sequencing data into 154 high-quality metagenomic assembled genomes (MAGs) in the fecal samples, enabling us to examine compositional changes at the strain level. Notably, we observed marked alterations in the human gut microbiota based on Bray-Curtis dissimilarity at the MAG level from day 0 to day 28 in both *B. lon_*8 and *B. lon_*1 group (R^2^=0.0608, p<0.05 in *B. lon_*8 group; R^2^=0.0563, p<0.05 in *B. lon_*1 group; PERMANOVA, **Figure 4A and Figure S6C**). Accordingly, the temporal change of *abfA* cluster abundance for a host did not correlate with the temporal Bray-Curtis dissimilarity between baseline and 28 days at the genus (r=-0.054, p=0.74, Spearman’s rank correlation, **Figure 4B**) or the species level (r=0.066, p=0.68, Spearman’s rank correlation, **Figure 4C**), whereas positively correlated with that at the strain level (r=0.55, p=0.00031; Spearman’s rank correlation; **Figure 4D**). Apparently, *B. longum-*induced shifts in the *abfA-*cluster abundance were mainly attributed to the abundance change from responsive resident strains that do not necessarily have close taxonomic relationships. Alternatively, these responsive strains might belong to the same genus or species taxa yet behaved radically different from others in the same taxonomic unit in response to the elevation of *abfA* cluster or just presented in a quite lower abundance than others in the same taxon.

**Figure 4.**
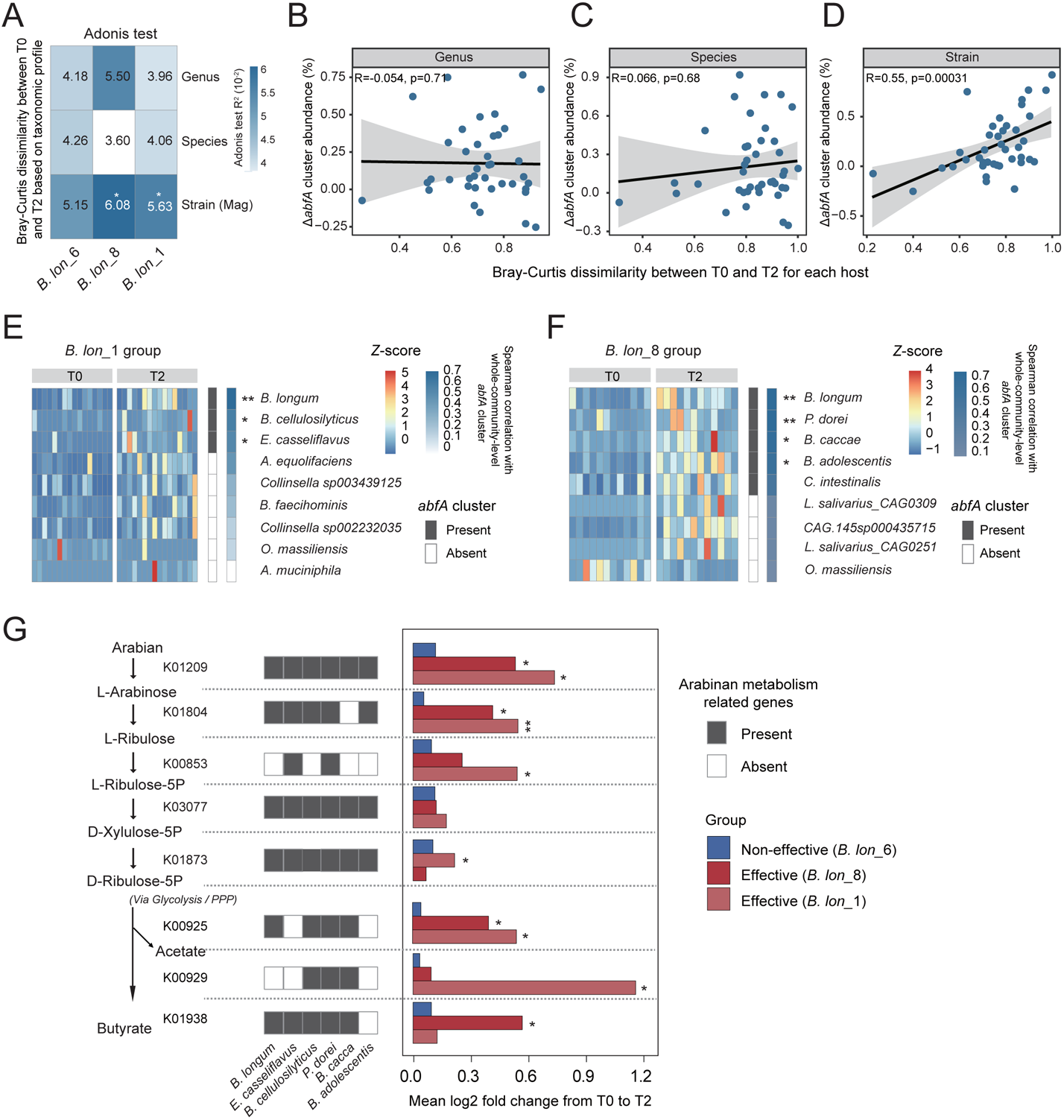
The shifts in gut resident microbiota responsive for elevated abundance of microbial *abfA* gene cluster after consumption of effective *B. longum*. (**A**) The variation of the intestinal microbiome of human subjects induced by *B. longum* with *abfA* gene cluster based on Bray-Curtis dissimilarity between T0 and T2 at different taxonomic profile. The heatmap shows R^2^ produced by the Adonis test. (**B-D**) The correlation between the change in *abfA* cluster abundance and the Bray-Curtis dissimilarity between baseline and 28 days at (**B**) genus, (**C**) species and (**D**) strains level, respectively. Linear model (LM) was used for creating a fit line for scatters. The error bands indicate the 95% confidence intervals. (**E**) The differentially abundant MAGs between the baseline and the 28 days after *B. lon_*1 consumption. The left panel, a heatmap shows the abundance of MAGs in each subject based on a zero-mean normalization (Z-score). The middle panel, a heatmap in black or white indicates if a strain carried the *abfA* cluster. The right panel, the Spearman correlation between the abundance of microbial strains and whole-community-level abundance of *abfA* cluster from baseline to the end point. (**F**) The differentially abundant MAGs between the baseline and the 28 days after *B. lon_*8 consumption. The left panel, a heatmap shows the abundance of MAGs in each subject based on zero-mean normalization (Z-score). The middle panel, a heatmap in black or white indicates if a strain carried the *abfA* gene cluster. The right panel, Spearman correlation between the microbial strain abundance and the whole-community-level abundance of *abfA* cluster from baseline to the end point. (**G**) Links between functional genes in the key arabinan-degradation pathway of gut microbiome and specific responsive gut microbes that exhibited changes in abundance with *B. longum* engraftment. The left panel, the black arrows indicate the key metabolic pathway of arabinan utilization in gut microbiome. The middle panel, the cells in black indicate the strains that carried the arabinan-metabolism-related genes. The right panel, the histograms showed that the change in abundance of arabinan-metabolism-related genes for effective (*B. lon_*6 and *B. lon_8*) and non-effective (*B. lon_*1) groups. *p<0.05, **p<0.01.

We next sought to detect gut microbes in response to the probiotic engraftment with a focus on MAGs. Again, we failed to identify any resident bacterial genera and species associated with the administration of effective *B. longum* in our cohort (p>0.05, Kruskal-Wallis test with Benjamini-Hochberg correction, **Figure S6A, Figure S6B** and **Table S4**), which was largely consistent with findings from many other studies ^49, 50^. Instead, 17 resident bacterial strains (MAGs) were identified exhibiting a significant change in abundance between baseline (T0) and day 28 (T2) in *B. lon_*1 or *B. lon_*8 group (p<0.05, Wilcoxon signed-rank test, **Figure 4E**, **Figure 4F** and **Table S4**). Out of these 17 resident strains, eight strains harbor the *abfA* cluster in their genomes. Out of these eight strains, three (*B. longum*, *B. cellulosilyticus* and *E. casseliflavus*) presented in subjects of the *B. lon_*1 group while five (*B. longum, P. dorei, B. caccae, B. adolescentis* and *C. intestinalis*) were identified in the *B. lon_*8 group (**Figure 4E**, **Figure 4F**, **Figure S6C and Figure S7**). We next performed correlation analysis within each group and found that all these eight *abfA-*cluster-carrying strains (expect for *C. intestinalis* in *B. lon_*8 group) positively correlated with the whole-community-level *abfA* cluster abundance (Spearman’s rank correlation, r>0.3, p<0.05, **Figure 4E, Figure 4F** and **Table S4**). These suggested that the administration of probiotic *B. longum* with *abfA* cluster populated the resident gut microbes that also can preferentially utilize arabinan.

To gain functional insight into the seven resident strains associated with overall fecal *abfA* cluster abundance, we next test if abundance change of these key strains enriched functional genes in the metabolic pathways related to arabinan degradation. We mapped metagenomic reads to Kyoto Encyclopedia of Genes and Genomes (KEGG) database to obtain the KO-term profile for metagenomic samples and compared KO abundances between time points in each of probiotic treatment groups (i.e., *B. lon_*1, *B. lon_*6 and *B. lon_*8). We observed that five KO terms (K01209, p<0.05; K01804, p<0.01; K00853, p<0.05; K09025, p<0.05 and K00929, p<0.05; **Figure 4**) in *B. lon_*1 group while five KO terms (K01209, p<0.05; K01804, p<0.05; K01873, p<0.05; K00925, p<0.05 and K01938, p<0.05; **Figure 4G** and **Table S5**) in *B. lon_*8 group, involving fermentation of L-arabinose to produce SCFAs, were drastically elevated at the end of the intervention compare to baseline. In addition, we found that these key resident bacterial strains harbored at least four out of eight genes in the arabinan metabolism.

From the molecular ecological point of view (**Figure 4G**), the supplementation of *abfA*-cluster-carrying *B. longum* strains promoted the hydrolysis of arabinan to produce a large amount of L-arabinose through catalyzing hydrolysis of L-arabinofuranosyl linkages found on the side chains of arabinan. The increased L-arabinose may selectively enrich resident bacterial strains resulting in the shifts in the abundance of fecal *abfA* cluster, as they could benefit each other through microbial cross-feeding of L-arabinose released by arabinan degradation. Since these resident bacterial strains commonly harbor an *abfA* cluster, the abundance change of these key strains typically accompanied with a marked increase in the whole-community-level *abfA* cluster abundance ^51, 52^.

### The elevated abundance of *abfA* cluster altered metabolic functions of gut microbiota

A variety of organic molecules produced by bacteria, including organic acids, pyrimidine derivatives, and tryptophan derivatives, have been shown to broadly influence host health ^35^. To gain insights into possible contributions of *abfA* cluster to the gut metabolic functions, we interrogated metagenomic and metabolomic features in stool samples from participants in our cohort at baseline and day 28. With the KEGG database, 14 KO terms altered significantly over time both in *B. lon_*1 and *B. lon_*8 group (p<0.05, Wilcoxon signed-rank test, **Figure 5A** and **Table S5**), while they remained unchanged in *B. lon_*6 group. These KOs involved in metabolic functions including arabinan metabolizing, acetate production, mannonate metabolizing, chenodeoxycholic-acid production and agmatine production (FDR<0.05 for all significant KO terms; K01442, log2(fc)=0.6096, p=0.0307; K00925, log2(fc)=0.6609, p=0.0407; K10012, log2(fc)=0.7642, p=0.0187; K01686, log2(fc)=-1.0620, p=0.0061; K01209, log2(fc)=0.7379, p=0.0338; **Figure 5A** and **Table S5**) in *B. lon_*1 group and (FDR<0.05 for all significant KO terms; K01442, log2(fc)=0.4012, p=0.0268; K00925, log2(fc)=0.5528, p=0.0225; K10012, log2(fc)=0.5610, p=0.0317; K01686, log2(fc)=-0.4786, p=0.0225; K01209, log2(fc)=0.5535, p=0.0374; **Figure 5A** and **Table S5**) in *B. lon_*8 group. We next tested whether the temporal change of functional KOs correlated with the shifts in fecal *abfA* cluster abundance. Interestingly, the temporal abundance changes in K10012 (*arnC*, L−arabinose transferase), (*rpmC*, large subunit ribosomal protein), K01804 (*araA*, L−arabinose isomerase), K01442 (*cbh*, choloylglycine hydrolase), K01209 (*abfA*, α−L−arabinofuranosidase) and K00925 (*ackA*, acetate kinase) were highly concordant with shifts in the overall *abfA* cluster abundance (Spearman’s rank correlation, r>0.3, p<0.05, **Figure 5A** and **Table S5**). Inversely, we also sought to probe what resident microbes primarily contributed to the overall abundance of a responsive functional gene (KO) in the gut. Expectedly, for example, *ackA* (K00925) positively correlated with *B. longum* and *B. adolescentis* in abundance (Spearman’s rank correlation, r>0.3, p<0.05, **Table S5**), which were previously reported to carry the *abfA* cluster in the genome.

**Figure 5.**
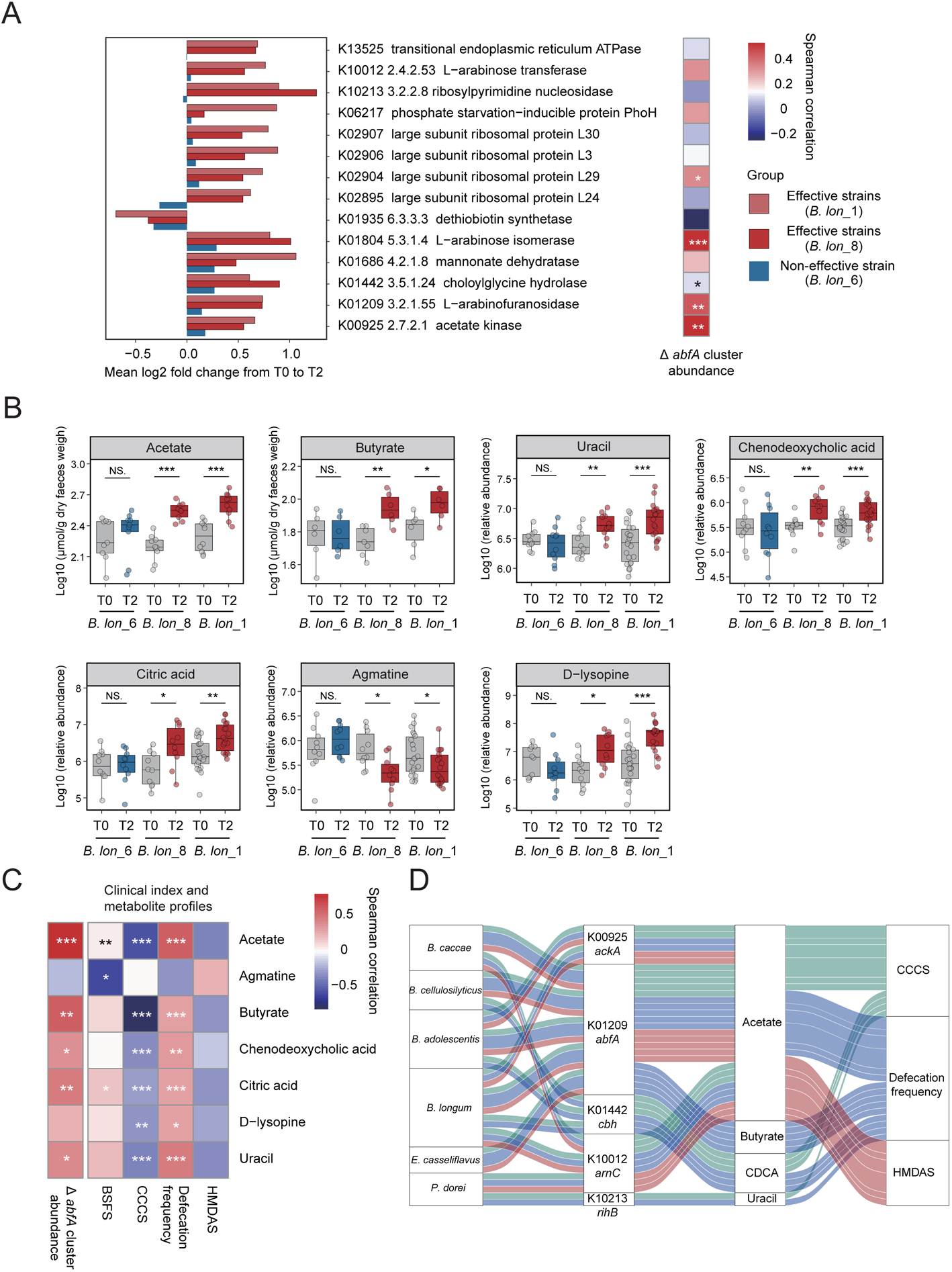
*B. longum* with *abf*A cluster alleviated constipation by modulating the indigenous gut microbial metabolic pathways and enhancing production of microbial beneficial metabolites. (**A**) The significant difference in the abundance of gut metagenome KO terms between the subjects on baseline and 28 days after probiotic consumption. The heatmap shows that Spearman correlation between the temporal change of KO modules and *abfA-*cluster abundance in gut microbiome. (**B**) The intestinal microbial metabolites that differed significantly between baseline and 28 days after probiotic consumption determined with LC/MS or GC/MS (N=10, 10, and 20 metabolite profiles for *B. lon_*6, *B. lon_*8, and *B. lon_*1, respectively). The p value from the linear mixed-effect models based on log10-transformed data were corrected with FDR. (**C**) Spearman correlation across the intestinal microbial metabolites, clinical index and fecal microbial *abfA* gene cluster abundance. (**D**) Sankey diagram showing the inferred the elaborate interplays among the microbial strains, microbial functional genes, fecal metabolites and clinical index with spearman correlation > 0.3. *p<0.05, **p<0.01, ***p<0.001 was determined by Wilcoxon signed-rank test (**B**), Data is presented with mean±SD.

To further explore the *abfA*-cluster-mediated changes in the fecal microbial metabolites, we first determined the concentration of fecal short-chain fatty acids (SCFAs) using gas chromatography-mass spectrometry (GC-MS). SCFAs are the major end products of carbohydrate metabolism in *Bifidobacteria* and other bacteria ^53^ and have been well documented to improve gut motility by interacting with GPR41 and GPR43, or act directly on colonic smooth muscle ^54–57^. Here, two major SCFAs (butyrate and acetate) produced by gut bacteria drastically increased after the 4-week treatment in both *B. lon_*1 and *B. lon_*8 group (Wilcoxon signed-rank test, p<0.05, **Figure 5B** and **Table S6**), whereas they didn’t change over time in *B. lon_*6 group. Next, we employed an untargeted metabolomics approach (LC-MS) to identify other fecal metabolic changes responsive for enriching *abfA* cluster abundance by probiotic administration. From the LC-MS profiles, uracil (p<0.01, *B. lon_*8 group; p<0.001, *B. lon_*1 group), chenodeoxycholic acid (p<0.01, *B. lon_*8 group; p<0.001, *B. lon_*1 group), citric acid (p<0.05, *B. lon_*8 group; p<0.01, *B. lon_*1 group) and D−lysopine (p<0.05, *B. lon_*8 group; p<0.001, *B. lon_*1 group) were enriched, while agmatine (p<0.05, *B. lon_*8 group; p<0.05, *B. lon_*1 group) was depleted in stool samples from patients at day 28 compared to baseline (Wilcoxon signed-rank test, **Figure 5B** and **Table S6**). We then intended to compare the within-host change of *abfA* cluster abundance to the those of the fecal metabolites, revealing the potential concordance between ecological and metabolic changes over time in the gut. Here, we observed the temporal changes in *abfA* cluster abundance in the gut microbial community positively correlated with those in acetate, butyrate, chenodeoxycholic acid, citric acid and uracil in feces (Spearman’s rank correlation, r>0.3, p<0.05, **Figure 5C**). Chenodeoxycholic acid was reported to increase stool frequency, decrease in stool consistency and increase the ease of stool passage via the membrane-bound bile acid GPBA receptor (TGR5) on enterocytes ^58–60^. The increased production of microbiota-induced uracil was associated with a decrease in IBS-C disease activity ^35^. Our study also supported those findings. The elevated production of acetate, butyrate, chenodeoxycholic acid, citric acid and uracil in the gut within each host correlated with the increase in defecation frequency (Spearman’s rank correlation, r>0.3, p<0.05, **Figure 5C**) as well as a decrease in CCCS (Spearman’s rank correlation, r>0.3, p<0.05, **Figure 5C**). Interestingly, at baseline, we also observed the significant correlations between these gut microbial metabolites and the between-host difference in FC symptoms (i.e., defecation frequency, and CCCS). (**Figure S6D**).

Finally, we integrated the multi-omics analysis results collected from the mouse model and human study to infer the interplays among gut microbes, metabolites and host phenotypes (**Figure 5D**). Overall, the administration of *B. longum* strains carrying *abfA* cluster enriched specific members that can efficiently utilize arabinan in the resident microbial community, affecting functional potentials and metabolite profiles in the gut microbiota. For instance, two functional genes (i.e., K01209 and K10012) derived from *B. longum*, *B. caccae* and *P. dorei* greatly contributed to the increased defecation frequency by elevating production of SCFAs (acetate and butyrate), which can increase the GI motility in either mice or humans ^55, 61^. Such links between acetate or butyrate and defecation frequency were also evident in our human cohort. The *cbh* (K01442) from *B. longum*, *B. adolescentis* and *B. cellulosilyticus* may contribute to gut transit time by affecting the concentration of fecal chenodeoxycholic acid. It was reported that chenodeoxycholic acid can accelerate colonic transit and improves bowel movement in patients with irritable bowel syndrome with constipation (IBS-C) and was deconjugated from their glycine or taurine conjugate by microbial bile-salt hydrolases in the host gut ^59, 62^. The enhanced production of uracil was mainly contributed by *rihB* (K10213) encoded by *P. dorei* and *E. casseliflavus* which correlated with CCCS score.

### Fecal microbiota transplantation experiment verified the protective role of *abfA* **cluster against FC**

To probe the causal role of gut microbial *abfA* cluster against FC, we next conduced a human-to-mouse fecal microbiota transplantation (FMT) experiment. The fecal samples from healthy human donors (N=6) with either enriched *abfA* cluster (mean relative abundance=0.926%, N=3) or depleted *abfA* cluster (mean relative abundance=0.031%, N=3) in the microbial community was transplanted to two groups of GF mice with loperamide-induced constipation (**Figure 6A**). These two groups of healthy fecal samples had a 29.86-fold difference averagely in the abundance of *abfA* cluster in the microbiome. GF mice transplanted with *abfA*-cluster- abundant fecal microbiota displayed overt constipation alleviation based on two key symptoms of FC. The whole gut transit time of GF recipients with *abfA-*cluster- abundant microbiota decreased by 50 mins, while their fecal output increased by 2 pallets/hr (**Figure 6B and Figure 6C**). No phenotypic changes were evident in GF mice receiving the fecal microbiota from healthy individuals with a lower abundance of *abfA* cluster (**Figure 6B and Figure 6C**). Therefore, we further demonstrated the causal role of *abfA* cluster of gut microbiota in the FC development and clinical treatment. To be noted, it also strongly suggested that the empiric administration of probiotics such as *B. longum* with no prior knowledge on the presence or absence of *abfA* cluster in the genome wouldn’t result in any effective alleviation of FC.

**Figure 6.**
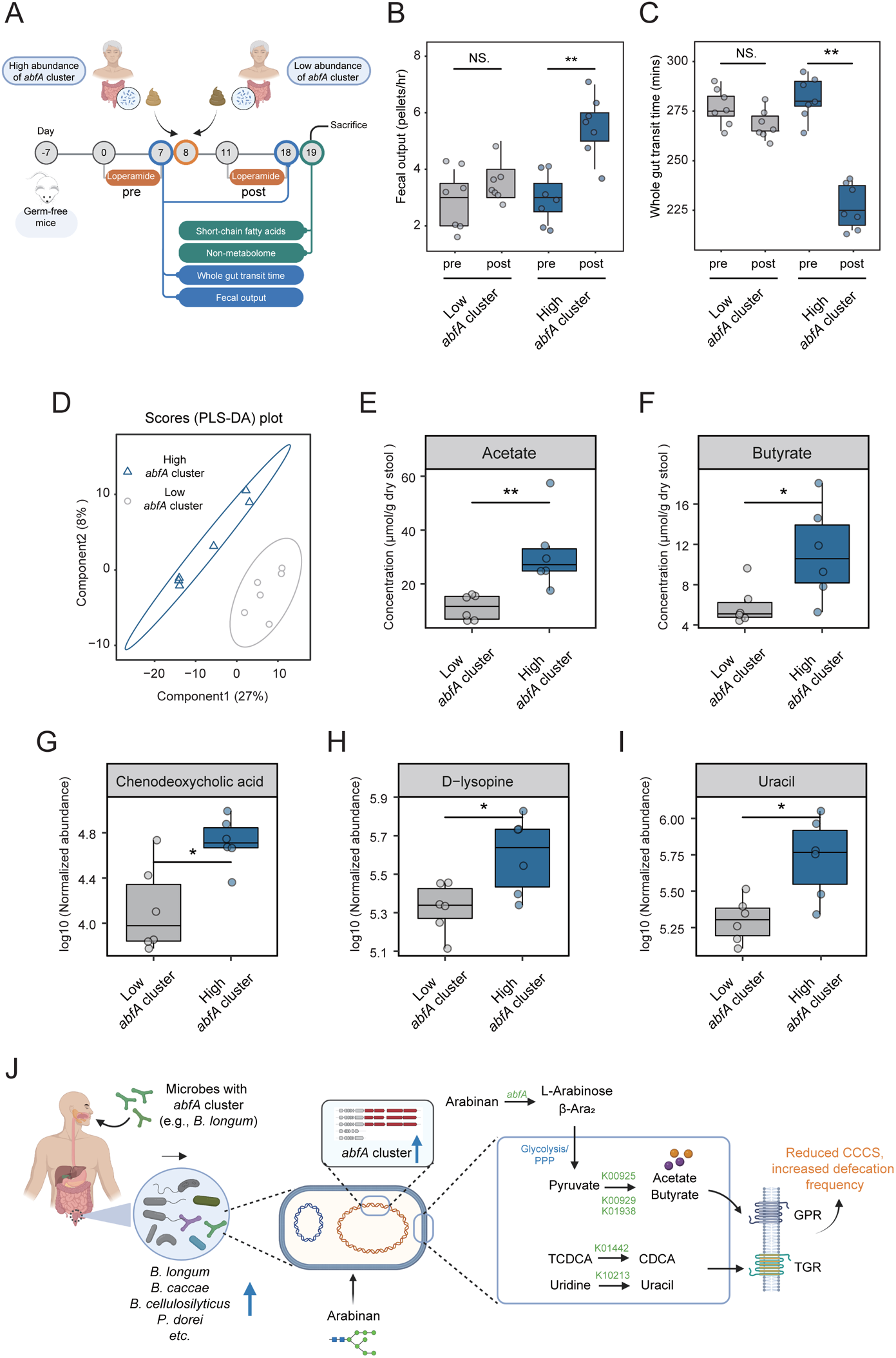
Transplantation of fecal microbiota with a high abundance of *abf*A gene cluster alleviated constipation induced by loperamide in GF mice. (**A**) Experimental design of GF mice by receiving fecal microbiota transplantation trial for further *abfA* gene functional verification. (**B-C**) Change in fecal output (**B**) and Whole gut transit time (**B**) in GF mice translated with high or low abundance of *abfA* gene cluster feces from human feces. (**D**) PLS-DA on metabolome data from mouse faeces after inoculation of high or low abundance of *abfA* gene cluster feces into germ-free mice (N=10). Proportions of the first components (Component1) and second components (Component2) are 27% and 8%, respectively. (**E-F**) Concentrations of the acetate (**E**) and butyrate (**F**) in the feces from each group at day 19 described in **Figure 6A**, determined with GC/MS. (**G-I**) Normalized abundance of metabolome profiles in the feces from each group at day 19 described in **Figure 6A**. N=10 mice per group. In all boxplots, the central line represents the median, box edges show the 25th and 75th percentiles, and whiskers extend to 1.5× the interquartile range. (**J**) Critical role of gut microbial with *abfA* cluster in alleviating functional constipation. *p[<0.05, **p<0.01, ***p[<0.001 as determined by Wilcoxon sum-rank test (**B and C**) or Student’s *t* test (**E-F and G-I**). Data is presented with mean±SD.

Next, we performed metabolomic analysis of feces from mice administered with *abfA*-cluster-enriched fecal microbiota (effective) and *abfA*-cluster-depleted fecal microbiota (non-effective). A striking difference in fecal metabolomic profiles between GF recipients from two treatment groups was observed (**Figure 6D and Table S6**) using PLS-DA. Furthermore, the majority of differentially abundant metabolites between two groups were largely consistent with those we identified in the human data. For example, acetate, butyrate, chenodeoxycholic acid, D-lysopine and uracil were over-abundant in the FMT-effective group of mice (**Figures 6E-6I and Table S7**). These data further demonstrated that this newly identified *abfA* gene cluster derived from either native gut microbiota or exogenous probiotics plays a key protective role against FC through the efficient utilization of plant polysaccharides (such as arabinan) that can’t be utilized by normal gut residents and simulating the production of a series of beneficial microbial metabolites improving the GI motility.

## DISCUSSION

Genetic factors governing the unique metabolic capability of probiotics should be primarily considered for screening probiotics or inferring their treatment efficacy against diseases. Conventionally, *Bifidobacterium, Lacticaseibacillus*, *Lactiplantibacillus*, etc. are major “probiotic” taxonomic groups. Strains from these taxa were empirically prescribed for treating FC due to the fact that the decreased abundance of *Bifidobacterium, Lacticaseibacillus*, *Lactiplantibacillus,* etc. was observed in patients with FC in the previous metagenomics studies. We here argue that only one single gene cluster can determine or stratify the key functional traits of probiotic strains (e.g., *B. longum*) that were taxonomically identical or genetically highly close to each other. Consistent with our findings, previous studies also demonstrated that a single gene or gene cluster can affect the cell surface composition or metabolite profile of a probiotic strain *in vivo*, which associated with particular physiological functions (i.e., bacterial adhesion ability, acid tolerance, and bile salt tolerance, etc.) ^17, 63, 64^. *Bifidobacteria BAtg* gene cluster encoding an ATP-binding-cassette-type carbohydrate transporter contributes to protecting mice from death induced by *E. coli* O157:H7 through the increased production of acetate by metabolizing fructose in the distal colon ^17^. Another study reported that *Bifidobacteria FL-SBP* gene cluster encoding putative ABC transporter SBP for fucosyllactose benefit infant by facilitating acetate production and limiting the abundance of *Enterobacteriaceae* ^64^. This motivated us to identify key genetic factor(s) determining the probiotic *Bifidobacteria* efficacy on improving the gut motility. In our previous studies, “effective” probiotic strains were often identified from a limited pool of candidate strains ^65–67^. A larger-scale genetic screen was achieved by establishing a comprehensive strain library in combination of paired genome sequence data. Next, we unprecedentedly established the causal link between a strain-specific genetic variation (*abfA* cluster) to the key functional difference of probiotic *B. longum* of our interest in the model organisms including mice and humans and provided mechanistic and ecological insights into how such a single gene cluster can affect the gut motility of hosts through arabinan metabolism.

The *abfA* cluster enables a highly efficient utilization of a significant portion of plant-based dietary fiber (arabinan) and facilitates the stable production of intestinal beneficial metabolites associated with the bowel movement. This *abfA* cluster encodes four linked genes all annotated as α-L-arabinofuranosidase, a glycoside hydrolase responsible for breakdown of bonds between sugars in arabinan, a common constituent of pectic polysaccharides ^68^. Arabinan is also known as the indigestible fiber for humans, or a poorly accessible source of nutrients to be utilized by normal gut microbiota. Yet, a mechanistic link between gut microbial arabinan metabolism and FC amelioration was largely underexplored. Here, we intended to elucidate the underlying mechanisms via the longitudinal multi-omics analysis of fecal samples from both mouse and human individuals receiving *B. longum* with and without *abfA* cluster. Due to the “priority effect”, the engrafted microbes first need to establish themselves in the gut ecosystem and outcompete other residents by creating their unique metabolic niches. As shown in **Figure 6J**, the *abfA* cluster confers effective strains (*B. longum*) a unique utilization ability of arabinan that was poorly utilized by many gut residents, which benefit their adaptation and colonization to the host gut. Based on a stable engraftment, probiotics can further functionally modulate gut microbiota in FC. In the mouse model, the arabinan-utilization capability of effective *B. longum* strains correlated with a high production of acetate in the gut, while other *abfA*-cluster-carrying residents promoted the production of butyrate or propionate. In the human cohort, we further recapitulated the observation that probiotic engraftment supported the growth of resident microbes with similar metabolic capability in utilizing plant polysaccharide. Interestingly, these microbial residents increased the production of a wider array of beneficial metabolites in the gut microbiota including acetate, butyrate, chenodeoxycholic acid and uracil. Acetate and butyrate are two well-known major SCFAs, end products of carbohydrate metabolism in *Bifidobacteria* and *Bacteroides* ^53^, which have been often implicated in improving intestinal motility by interacting with G-protein-coupled receptor 41 (GPR41) and G-protein-coupled receptor 43 (GPR43), or directly acting on colonic smooth muscle ^54–57^. Those findings, together with *Bifidobacteria* and *Bacteroides* harboring the SCFA-producing genes (K00925 acetate kinase) and arabinose-degrading genes (K10012 L−arabinose transferase and K01804 L−arabinose isomerase), supports our notion that *B. longum* with *abfA* cluster alleviated FC by establishing specific metabolic niche in the host gut and further enhancing the SCFA-production pathways. Besides, the abundance of *Bifidobacteria* and *Bacteroides* were also associated with the bile acid level (i.e., chenodeoxycholic acid) in the gut ^35, 69^. In our study, the increased abundance of these two taxa slightly correlated with the elevated level of choloylglycine hydrolase (K01442), mainly contributing to the enrichment of bile-acid metabolic pathway in the gut microbiota. For patients with irritable bowel syndrome with constipation (IBS- C), it is reported that treatment with chenodeoxycholic acid can increase stool frequency, decrease stool consistency and improve ease of stool passage by acting on the membrane-bound G protein-coupled bile acid receptor (e.g., TGR5) on enterocytes ^58–60^. The uracil metabolism was also enriched in the gut microbiota after the engraftment of *B. longum* carrying *abfA* cluster, which was associated with the increase of abundance of *P. dore* strain harboring *richB* (K10213), a gene involved in uracil metabolism. Consistent with our findings, a previous study reported the link between the increase of microbiota-induced uracil levels and the decrease of IBS-C disease activity ^35^. Notably, in the proximal part of colon, the glucose supply is often sufficient for the microbial production of SCFAs, bile acids and uracil, whereas, in the distal colon, the glucose supply should be exhausted. In this circumstance, only microbes equipped with this *abfA* cluster can stably proliferate and produce beneficial metabolites aforementioned. Taken together, our results clearly explain why only a probiotic strain carrying the *abfA* cluster can stably modulate the gut microenvironment and increase the GI motility.

Microbial biomarker discovery is also one key goal in human gut microbiome studies, boosting the development of rapid, non-invasive diagnostic or prognostic approaches in the clinical practice ^70–72^. We, for the first time, demonstrated that the *abfA* cluster abundance of fecal microbiome can be developed as a highly accurate biomarker to predict the symptom severity of FC and potentially a novel assessment method for broad GI issues. Similarly, Wirbel et al. reported that the copy number of the gut microbial gene *baiF* (encoding metabolic pathways producing secondary bile acids including 7α-dehydroxylation) clearly distinguished CRC patients from health controls (p=0.001) with an AUROC of 0.77, suggesting the feasibility of using a single gene marker in the gut microbiome for prediction of GI health status ^71^. Our FMT studies further supported the causal role of gut microbial *abfA* cluster for FC, highlighting the importance of its prognostic value in the FC treatment. On the other hand, this gene cluster can serve as a tangible genetic marker to predict the effectiveness of probiotics against FC via a cost-effective PCR test. Last but not least, it was found to be in a lower abundance in patients with T2D, CRC, ACVD, suggesting its essential roles in the host gut health. Those tested chronic diseases or gastrointestinal diseases all likely or at least partially resulted from a lack of high-fiber diet or a deficiency in the key functions of gut microbiome for sufficiently degrading dietary fibers, associated with a limited production of beneficial SCFAs (e.g., acetate and butyrate) modulating gut dysbiosis. Qin et al. reported that a decrease in the level of bacterial butyrate biosynthesis in the gut microbiota of T2D patients ^44^. In ACVD patients, the butyrate-producing bacteria including *R. intestinalis* and *F. prausnitzii* were underrepresented as well ^48^. Feng et al. reported that a diet containing a low fiber relative to meats would lead to the outgrowth of putrefactive bacteria, promoting colorectal carcinoma ^41^. Together, the *abfA* cluster responsible for degrading a unique plant polysaccharide may not only a causal factor to FC but also contribute to the diagnosis or prediction of T2D, ACVD, and CRC, etc.

Collectively, this study identified and systematically characterized a key genetic factor responsible for arabinan utilization that directly addressed one of most critical challenges in the probiotic field, that is, a widespread yet unknown variability in treatment efficacy across probiotic strains. Technically speaking, our proof-of-concept study also established generalizable principles for rational development of colonizable, functional probiotics with a consistent and persistent treatment efficacy in multiple model organisms. Essentially, the equipment with *abfA* cluster in the genome offered ingested probiotics fitness advantages over other gut residents by creating metabolic niches for stable production of beneficial metabolites against FC. Furthermore, it is so prevalent in the gut microbiota that can be developed as a simple yet powerful functional marker for FC and other GI-related diseases. On the other hand, we also underscored the importance of examining and rescuing the silent genetic deficits in the gut microbiota of humans for digesting dietary fiber (such as *abfA* cluster) in combating many other fundamentally non-addressable chronic diseases or gut microbial dysbiosis.

## STAR METHODS

Detailed methods are provided in the online version of this paper and include the following:

- KEY RESOURCES TABLE
- RESOURCE AVAILABILITY

◦ Lead contact
◦ Materials availability
◦ Data and code availability
- EXPERIMENTAL MODEL AND SUBJECT DETAILS

◦ Animal experiments
◦ Human volunteers
- METHOD DETAILS

◦ Bacterial strain isolation from fecal samples and genome sequencing
◦ Phylogenetic analysis and functional annotation for isolated *B. longum* strains
◦ *In vitro* growth assay of isolated bacterial strains
◦ *In vitro* transcriptional activity of key gene clusters
◦ Shotgun metagenomic data analysis
◦ Metabolomics analysis with LC-MS
◦ The quantification SCFAs with GC-MS
◦ Measurement of fecal pellet water content of mice
◦ Determination of whole gut transit time of mice
◦ Determination of small intestinal transit and blood sample collection
◦ Measurement of gastrointestinal hormones related to constipation
- QUANTIFICATION AND STATISTICAL ANALYSIS

### KEY RESOURCES TABLE

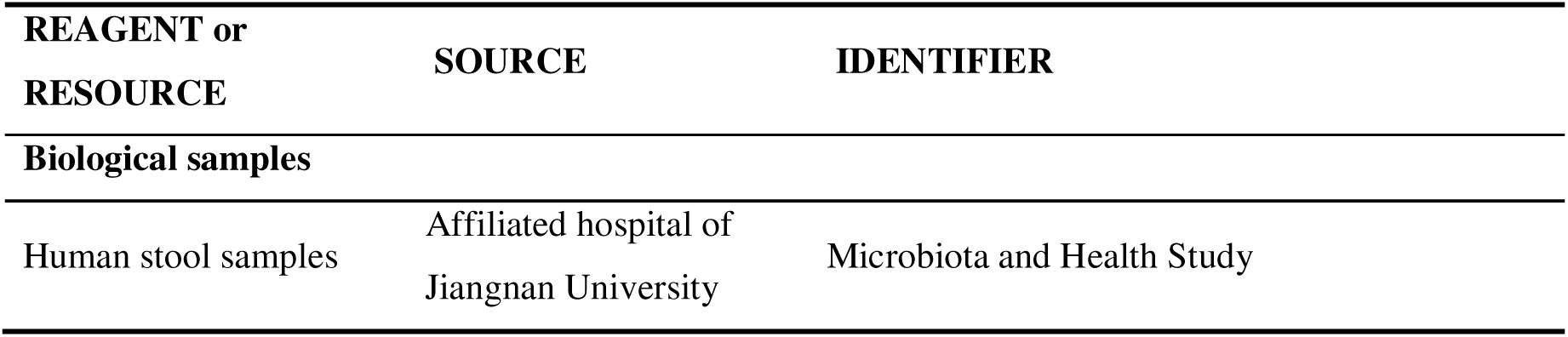
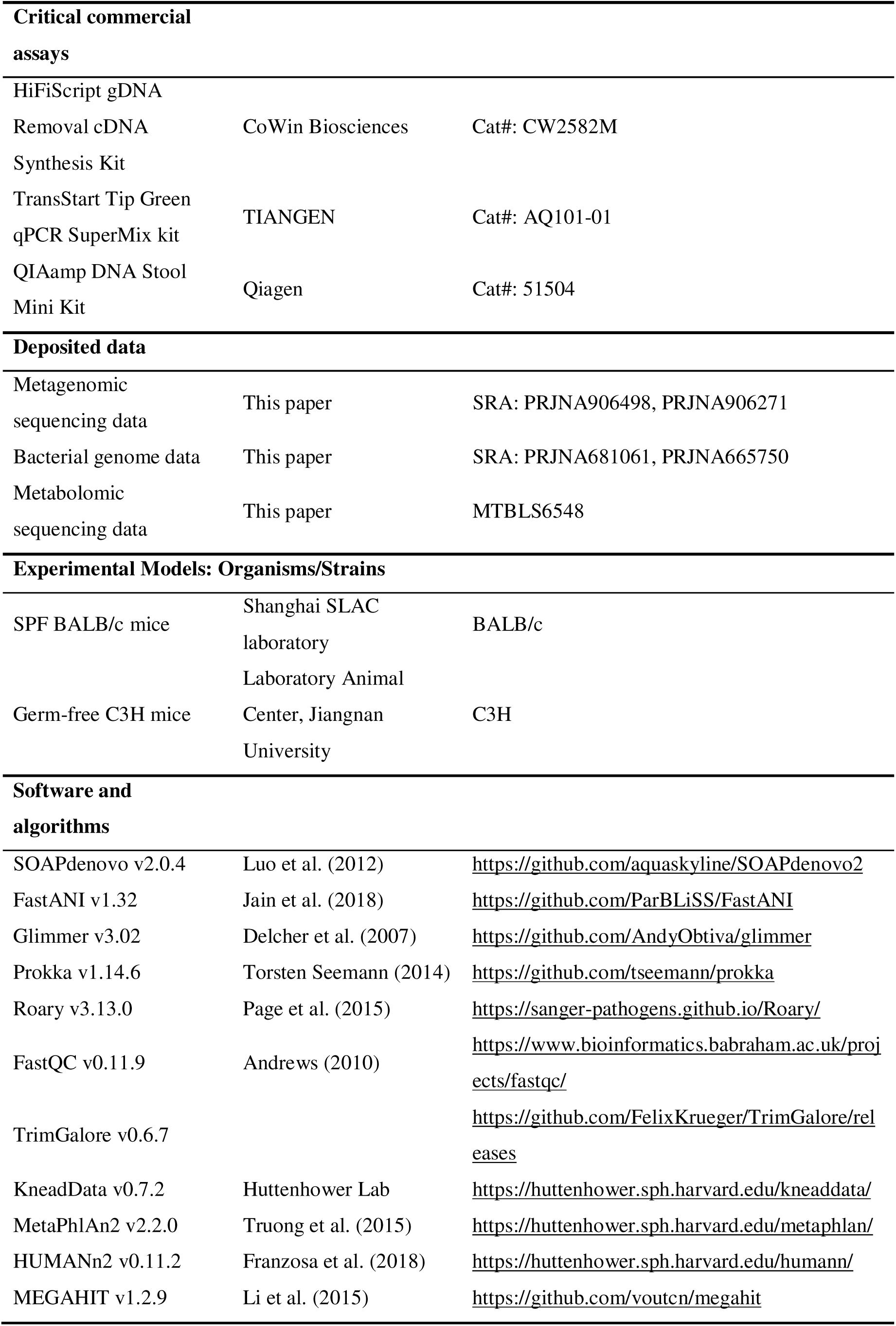
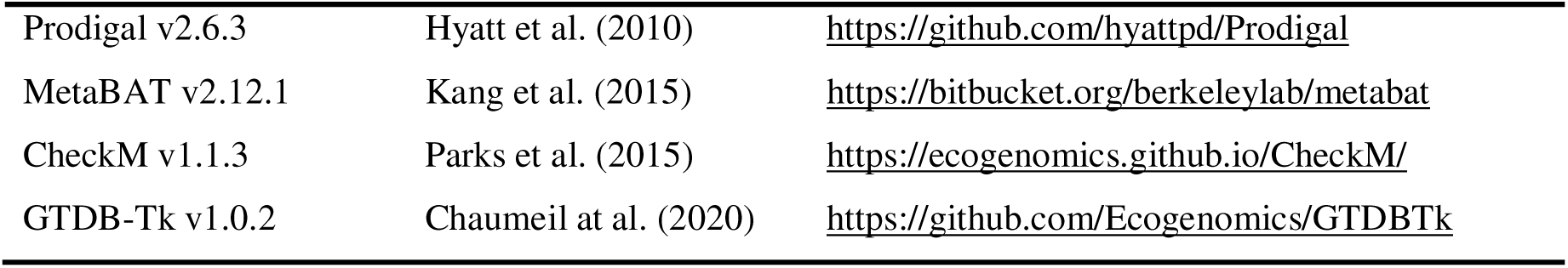

### RESOURCE AVAILABILITY

#### Lead contact

Further information and requests for resources should be directed to and will be fulfilled by the lead contact, Qixiao Zhai.

#### Materials availability

This study did not generate new unique reagents.

#### Data and code availability

- Metagenomic data are publicly available in NCBI Sequence Read Archive under BioProject: PRJNA906498, PRJNA906271. Metabolomic data are deposited at MetaboLights repository: MTBLS6548 (www.ebi.ac.uk/metabolights/MTBLS6548). Clinical data are available upon request from Qixiao Zhai. Bacteria isolates were deposited in the Culture Collection of Food Microorganisms (CCFM), Jiangnan University (Wuxi, China) and obtained genome sequencing data are publicly available in NCBI Sequence Read Archive under BioProject: PRJNA681061 and PRJNA665750.
- Analysis software including quality control, taxonomic, and functional profiling tools are publicly available and referenced as appropriate. Code used for metagenome and metabolome data analysis is available at https://github.com/ProC20/FC.git.
- Any additional information required to reanalyze the data reported in this work paper is available from the lead contact upon request.

### EXPERIMENTAL MODEL AND SUBJECT DETAILS

#### Animal experiments

Male SPF BALB/c mice (8 weeks old, 26–28 g, Shanghai Slack Experimental Animal Co., Ltd., China) were used for the animal experiments. Mice were adapted to standard environmental conditions (temperature, 25±2°C; humidity, 55%±5%) with a 12-hr light-dark cycle for one week. The mice were provided with standard commercial mouse food and sterile water ad libitum during the experiment. The animal experimental protocols were approved by the Ethics Committee of Jiangnan University, Wuxi, China (JN. No20180615b0950901[164], JN.No20190315b0940423[28], JN.No20190315b0940423[28], JN.No20200710b0700820[158], JN.No20210630b0400820[221]), and were performed in accordance with the guidelines established by the European Community (Directive 2010/63/EU).

#### Animal Experiment 1

To evaluate the effects of *B. longum* strains on loperamide-induced constipation, 80 mice were randomly divided into eight groups. All animals were fasted overnight (approximately 12 hr) before the first experiment while water was not restricted. Subsequently, the mice were administered with a gavage dose of loperamide (10 mg kg^-1^) to induce constipation once per day for 14 continuous days. At 1 hr later administration, all mice were treated with four *B. longum* strains (*B. lon_*70, *B. lon_*4, *B. lon_*8, *B. lon_*6 and *B. lon_*39) at a sufficient dose (1.0 × 10^9^ CFU per day) by the same method, or sterilized PBS (0.01M, pH 7.4) as a control group or a single dose of phenolphthalein (7 mg kg^-1^) as a positive drug control group. Moreover, the mice were given only sterilized PBS (0.01M, pH 7.4) without loperamide administration as a normal group.

#### Animal Experiment 2

To assess the effects of specific *B. longum* strains with two arabinan-degrading gene clusters on loperamide-induced constipation in mice, the SPF BALB/c mice (N=6∼8 biologically independent animals per group) were administered with loperamide and treated with eight *B. longum* strains (*B. lon*_56 and *B. lon_*63 with only *abfA* cluster, *B. lon*_28 and *B. lon*_58 with only *hypBA cluster* , *B. lon*_24 and *B. lon*_23 with *abfA* cluster and *hypBA* cluster, or *B. lon_*64 and *B. lon*_30 without *abfA* cluster and *hypBA* cluster) or sterilized PBS (0.01M, pH 7.4) as a control group at 1 hr after loperamide administration or sterilized PBS (0.01M, pH 7.4) as a normal group without loperamide administration, respectively.

#### Animal Experiment 3

To further assess the effects of specific *B. longum* strains with *abfA* cluster on loperamide-induced constipation in mice, the SPF BALB/c mice (N=6∼8 biologically independent animals per group) were administered with loperamide and treated with *B. lon_*8, *B. lon_*6, arabinan + *B. lon_*8, arabinan + *B. lon_*6 or arabinan alone.

#### Animal Experiment 4

To assess the efficacy of more gut microbial strains (*B. longum*, *B. breve*, *P. pentosaceus* and *B. nordii*) with or without *abfA* cluster on loperamide-induced constipation in mice, the SPF BALB/c mice (N=6∼8 biologically independent animals per group) were administered with loperamide and treated with *B. lon*_3 with *abfA* cluster and *B. lon_*1 without *abfA* cluster, *B. bre*_27 with *abfA* cluster, *B. bre*_41 without *abfA* cluster, *P. pen_*16 with *abfA* cluster, *P. pen_*35 without *abfA* cluster, *B. nor*_12 with *abfA* cluster, *B. nor*_19 without *abfA* cluster or sterilized PBS (0.01M, pH 7.4) as a control group at 1 hr after loperamide administration or sterilized PBS (0.01M, pH 7.4) as a normal group without loperamide administration, respectively.

#### Transplantation of human fecal microbiota to germ-free (GF) mice

Sterile phosphate buffered saline (PBS) is pre-reduced by placing it in an anaerobic chamber overnight to remove oxygen. 10% (w/v) freshly thawed human feces were suspended in pre-reduced PBS under anaerobic conditions, and the fecal slurry was rotated for 5 mins ^73^. Centrifuge at 1000 g for 3 mins, and then filter with a 100-μm sterile filter to separate the suspended bacteria from the fiber material. Under anaerobic conditions, the resulting slurry (300 μL) was placed in a single cryotube. The male or female C3H GF mice aged at 8-9 weeks were randomly divided into two groups (N=10 animals per group; male: female = 5:5), followed by an oral gavage of loperamide at the dose of 10 mg kg^-1^ (maximum 250 μL). In order to avoid the co-housing of male and female mice, male or female mice (N=5) in each group were co-housed in a separate ventilated cage and had free access to autoclaved water and rodent feed. Every 7 days, the mice are placed in a clean cage equipped with autoclaved mattresses, sheds, nests, water and food. After the experiment, mice were euthanized with CO_2_, followed by cervical dislocation ^74^.

#### Human volunteers

Participants aged between 65-105 years who submitted the written informed consent prior to the study were invited to undergo the eligibility-assessment procedure. Subjects meeting the following criteria were included: *i*) fulfilled Rome IV diagnostic criteria for FC ^75^, including self reported stool frequency of 3 or less bowel movements per week, and self reported stool consistency of type 1 4 based on the Bristol Stool Form Scale; *ii*) body mass index (BMI) ranged between 18.5 29.9 kg/m^2^. A total of 10 healthy human subjects matched age and BMI with FC patients were recruited in this study.

Exclusion criteria included: *i*) volunteers with prior history of chronic gastrointestinal or nervous-system disease, or other conditions that may affect intestinal motility, *ii*) experienced major abdominal surgery (especially gastrointestinal surgery), *iii*) drug use related to digestive tract diseases within 28 days (anti-spasmodic drugs, anti-diarrhea drugs or laxatives), *iv*) moderate or severe anorectal problems, *v*) allergy or lactose intolerance to the ingredients of experimental probiotic products, *vi*) antibiotic use within the past 28 days vii) bleeding risk or taking medications that increase the risk of bleeding.

The elderly participants were required to submit the questionnaire on the basic health information, dietary structure, living condition, under the appropriate assistance at the first visit. At each study visit, participants also filled the questionnaires for assessing the functional-constipation-related phenotypic changes throughout the study, including stool consistency using the Bristol Stool Form Scale (BSFS) ^76^, stool frequency assessed by a daily stool diary, Cleveland Clinical Constipation Score (CCCS) ^77^, Patient Assessment of Constipation Symptoms (PAC SYM) ^78^, a 7-day Bowel Diary, and Patient Assessment of Constipation Quality of Life (PAC[QoL) ^79^. Stool samples for each subject were collected using a home collection kit throughout the study. Sample tubes instantly preserved in frozen gel packs were returned within 24 hrs to the Department of gastroenterology, Affiliated Hospital of Jiangnan University, where samples were immediately stored at −80°C. Each subject in the healthy control group (N=10) only provided one stool sample using the same protocol. Clinical trials were approved by the Medical Ethics Committee of Affiliated Hospital of Jiangnan University (IEC2020061204 and IEC2020092903). Human tissue samples were obtained from individuals who provided informed consent. Complete clinical trial registration is deposited at Chinese Clinical Trial Registry (ChiCTR2000034415 and ChiCTR2100041925, http://www.chictr.org.cn/).

### METHOD DETAILS

#### Bacterial strain isolation from fecal samples and genome sequencing

A total of 354 fecal samples of Chinese residing in 17 provinces or municipalities were collected for isolating *B. longum* strains. One hundred and eighty-five *B. longum* strains were isolated from Chinese fecal samples and their whole genome was sequenced in this study. The strains were cultured in mMRS plates (supplemented with 0.05% [w/v] L-cysteine and 50 mg/mL mupirocin) in a Whitley DG250 anaerobic workstation (Don Whitley Scientific Limited, Shipley, UK) and incubated at 37°C for 48 hrs. Then a DNA extraction kit (OMEGA, Biotech, Doraville, GA, USA) was used to extract genomic DNA from the bacterial cultures. Genome sequencing was conducted by the Illumina HiSeq Platform, which generated 350-bp pair-end reads. SOAPdenovo v2.0.4 ^80^ was used to assemble the paired-end reads *de novo* into high quality sequences. GapCloser was used to fill in the gaps inside each scaffold and correct single base errors.

#### Phylogenetic analysis and functional annotation for isolated *B. longum* strains

The 185 *B. longum* genomes were re-annotated using Prokka v1.14.6 ^81^, and a pan-genome analysis was conducted via Roary v3.13.0 ^82^ to identify the core genes of these *B. longum*. The genome of *B. longum* JCM 1217 (NC_015067.1) was downloaded from NCBI and used as a reference genome. The CDSs of genes were predicted for each sequenced genome by using Glimmer v3.02 ^83^. To obtain functional annotation, the amino-acid sequences of predicted CDS were blasted (BLASTP) against the NCBI nr and COG database ^84, 85^ with the criterion of e-value < 1e-5, identity > 40% and length coverage of gene > 50%. Strain-specific functional genes or gene clusters potentially linked to probiotic phenotypes were next identified. The R package “gggenes” was used for drawing gene arrow maps of *abfA* and *hypBA* gene clusters specific to effective probiotic strains. The pair-wise ANI values across newly sequenced genomes were calculated using FastANI v1.32 ^86^.

#### *In vitro* growth assay of isolated bacterial strains

The strains were inoculated in sugar-restricted basal medium supplemented with 1.0% D-glucose and cultured at 37°C under anaerobic conditions until the optical density at 600 nm (OD_600_) reached 1.0. A 5% volume seed culture was added to basal medium containing 2.0% arabinan or 2.0% D-glucose. The wells of a 96-well plate were filled with these mixtures and sealed with Microseal B plate-sealing film (Bio-Rad, USA). The plate was maintained at 37°C, and the OD_600_ was monitored every 2 hrs using a Powerscan HT plate reader (Dainippon Sumitomo Pharma, Japan).

#### *In vitro* transcriptional activity of key gene clusters

*B. lon_*8 was cultured at 37°C under anaerobic conditions in sugar-restricted basal medium with either 2.0% arabinan or 2.0% D-glucose. Cells in the culture of strains (1 mL) were collected via centrifugation (8000 g, 10 mins) in an enzyme-free tube and then mixed with 200 μL of lysozyme solution. The mixtures were incubated at 37°C for 30 mins to break the cell walls. Cells without cell walls were collected via centrifugation (12000 g, 5 mins, 4 °C). Total RNA from each strain was extracted using the TRIzoI Plus RNA Purification Kit according to the manufacturer’s protocol (Invitrogen, Carlsbad, CA, USA). The RNA concentrations were measured using an RNA assay with a NanoDrop spectrophotometer (Thermo, Shanghai, China). The cDNA was synthesized in a 20 μL reaction using a HiFiScript gDNA Removal cDNA Synthesis Kit (CoWin Biosciences, Wuxi, Jiangsu, China). qRT-PCR was performed in a 96-well plate following the TransStart Tip Green qPCR SuperMix kit (Tiangen Biotech Co., Ltd., Beijing, China) instruction. The specific primers were designed using Primer5 software (CA, USA) and were synthesized by the Sangon Biotechnology Laboratory (Shanghai, China). The 16S rDNA was used as a reference gene ^87^. Each experiment was conducted with triplicates.

#### Fecal metagenomic sequencing

DNA extracted from the fecal samples using the QIAamp DNA Stool Mini Kit (Qiagen, Hilden, Germany), examined by 0.8% agarose gel electrophoresis, and the OD value of 260/280 was determined by spectrophotometry. All DNA samples were subjected to shotgun metagenomic sequencing by using a HiSeq 2500 (Illumina, CA, USA) with 150 bp paired-end reads in the forward and reverse directions by Novogene Company (Beijing, China). The quality of sequencing reads was controlled by FastQC v0.11.9 ^88^ and followed by trimming and removal of human sequences with KneadData v0.7.2.

#### Shotgun metagenomic data analysis

Quality-filtered metagenomes were taxonomically profiled using MetaPhlAn2 v2.2.0 ^89^. Functional analysis was performed using HUMAnN2 v0.11.2 ^90^ in the UniRef90 mode with default settings. Accordingly, we obtained the relative abundance profiles of intestinal microbial taxa, gene families and metabolic pathways for each metagenome. Next, metagenome-assembled genomes (MAGs) were constructed using a workflow that included genome assembly by MEGAHIT v1.2.9 ^91^, prediction of open reading frames by Prodigal v2.6.3 ^92^, genome binning by MetaBAT v2.12.1 ^93^, genome quality control by CheckM v1.1.3 ^94^ and followed by taxonomic identification with GTDB-Tk v1.0.2 ^95^.

#### Metabolomics analysis with LC-MS

For extraction, each fecal sample from each individual mouse (about 50 mg) was immediately quenched in liquid nitrogen and mixed with 400 pre-cooled mixture of ultra-pure water and methanol (1:4, v/v), and homogenized with high-throughput tissue homogenizer SCIENTZ-48 (SCIENTZ, Ningbo, Zhejiang, China) at −20 for 6 mins. Each sample then vortexed for 30 s, and incubated with low-temperature ultrasonic extraction for 30 mins (5, 40 KHz); the stored at −20 for 30 mins. The test tube containing the sample was the centrifuged at 1300 g at 4 [ for 15 mins to obtain the supernatant. Quality control (QC) samples were prepared by pooling an equal volume of reconstitution from every plasma sample and were analyzed after every 10 plasma samples. QC samples can monitor the system stability and reproducibility. Blank sample (80% methanol) was used for background subtraction and noise removal.

The LC-MS analysis of the metabolite extracts (supernatant) of stool samples was performed using a UltiMateU-3000 UPLC system (Thermo Fisher Scientific, MA, USA) coupled with a high-resolution Q-Exactive mass spectrometer in full scan mode (Thermo Fisher Scientific, MA, USA). A 2.1 × 100 mm reverse-phase Waters Acquity UPLC T3 column (Waters, MA, USA) was used for chromatographic separation at 35°C. The mobile phase consisted of 0.1% aqueous formic acid (A) and 0.5 mM ammonium acetate-acetonitrile (B) with a gradient elution: 0–1.0 mins (5%, B); 1.0– 10 mins (5–99%, B); 10–12 mins (99%, B) and 12–15 mins (5%, B). The flowrate was 0.3 mL min^−1^ and the injection volume was 2 ml. The MS system with a heated electrospray ionization (ESI) source was operated in the positive and negative ion modes with a full-scan range covered 151 to 2000 m/z and 70 to 1050 m/z, respectively. And the parameters were set as follows: 1.50 kV source voltage and 250°C capillary temperature. Capillary voltages for negative and positive ionization modes were set at 2500 and 3500 V, respectively.

MS data were processed by Compound Discovery 3.1 (Thermo Fisher Scientific, MA, USA) using untargeted metabolic workflow with default parameters, which included data extraction, background features filtered out, peak identification, deconvolution, alignment and integration, and metabolite identification assignment via mzCloud, HMDB, KEGG, and ChemSpider databases. The data processing filtrated out metabolic features that appeared < 50% of the QC samples and that > 30% RSD of the QC samples, and the mass tolerance was 5 ppm. Metabolic features that whose m/z ratios could not match the masses in the above databases were filtrated out. The processed data including peak area, retention time (RT), molecular weight (MW), and identified or unknown compound were integrated in an excel. Multivariate analysis was performed in SIMCA software (Umetrics, Umea, Sweden). Metabolic pathway analysis was performed in MetaboAnalyst 5.0 (https://www.metaboanalyst.ca/).

#### The quantification of SCFAs with GC-MS

A total of 0.2 g of each stool sample was resuspended in saturated NaCl solution (500 μL), where 20 μL of sulfuric acid (10%) was added for acidification. 800 μL of diethyl ether was then added into the sample to extract short-chain fatty acids. After shaking for 2[mins, the mixture was centrifuged at 14,000 g for 15 mins, followed by adding 0.25 g anhydrous Na_2_SO_4_ to remove inside water. Keep at −20°C for 30 mins. Transfer the supernatant into a fresh glass vial, for GC-MS analysis. GC-MS analysis was performed using a gas chromatography-mass spectrometry system (GCMS-QP2010 Ultra system, Shimadzu Corporation, Japan). The system utilized a Rtx-Wax column (30 m × 0.25 mm × 0.25 μm, Restek, Evry, France). A 1-μL aliquot of the analyte was injected in a split mode (10:1). Helium was used as the carrier gas, the front inlet purge flow was 2 mL/min, and the gas flow rate through the column was 1 mL/min. The initial temperature was kept at 100 °C for 1 min, then raised to 140 °C at the rate of 7.5 °C/min, followed by 60 °C/min to 200 °C, and hold at 200 °C for 3 mins. The injection, transfer line and ion source temperature were set at 240°C, 240°C and 220°C. The analytes were detected using the single scan mode (acetic, butyric, isovaleric and valeric acids: 60 m/z; propionic acid: 57[m/z; isobutyric acid: 43 m/z). The concentration of SCFAs was calculated by calculating responsive factors for each SCFA relative to 4-methyl-valeric acid using the injections of pure standards, expressed in μmol/g dry sample.

#### Measurement of fecal pellet water content of mice

The number of fecal pellets for each mouse was first recorded for every 30 mins in a 5-hr experiment for measurement of gut motility of mice. The average number of fecal pellets for each mouse per hr was measured for inter-individual comparisons. One fresh fecal pellet from each mouse was then collected in a separate sterile EP tube. After obtaining the wet weight, each sample was subject to freeze dryer for 48 hrs to obtain the dry weight. The fecal water content for one mouse was calculated as the difference between the wet and dry weight of this representative fecal pellet.

#### Determination of whole gut transit time of mice

At the endpoint (e.g., day 14) of each animal experiment, mice were fasted overnight while water was provided. Then, the mice in the administration group were given loperamide (10 mg kg^-1^ body weight), and mice in the control groups were treated with 0.9% saline. One hr later, all mice were given oral gavage of 250 μL of 0.5% activated charcoal solution in combination with gum powder which can relief the potential irritation of the mouse intestine caused by introducing activated charcoal. The mice were immediately transferred to a clean, empty individual cage and allowed to eat and drink at will. The whole gut transit time or defecation time was recorded as the time required for defecation of first toner-containing stool pallet, i.e., darkening stool.

#### Determination of small intestinal transit and blood sample collection

The gastrointestinal motility was measured using the method adopted from Verdú et al. with slight modifications ^96^. After 12-hr fasting, each mouse was given activated carbon solution by gavage aforementioned. Firstly, to measure blood-derived gastrointestinal hormones, the mice were anesthetized with light ether for blood sample collection. The whole blood drawn from each mouse was placed in the tube for nearly 2 hrs allowing the formation of a clot. The serum was then separated from the clot by centrifugation at 3000 g for 15 mins. Next, the abdomen of each mouse was opened, the entire small intestine (from the pylorus to the cecum) was carefully taken out and placed it on blotting paper. The distance moved by the activated charcoal and the total length of the small intestine were measured. The small intestinal transit of each mouse was calculated as the percentage of the moving distance of the activated charcoal powder relative to the total length of the small intestine.

#### Measurement of gastrointestinal hormones related to constipation

The concentration of four representative gastrointestinal hormones, namely motilin (MTL), gastrin (Gas), somatostatin (SS) and vasoactive intestinal peptide (VIP) in the mouse serum was quantified using ELISA kits (Senbeijia Biotechnology, Nanjing, China) according to the manufacturer’s instructions. Briefly, after addition of the four specific hormone antibodies, separately in each well, the serum was incubated for 30 mins at 37, following which HRP-Conjugate reagent was added and incubated for 30 mins at 37[. The TMB One-Step Substra te Reagent was then added, and the mixture was further incubated for 15 mins at 37 . The reaction was terminated following addition of the sulphuric acid solution (the blue color change to yellow color). Finally, the absorbance of the reaction mixture was read at 450 nm after adding stop solution and within 15 mins using a spectrophotometer (Multiskan GO, Thermo Fisher Scientific Oy Ratastie 2, FI-01620 Vantaa, Finland).

### QUANTIFICATION AND STATISTICAL ANALYSIS

All statistical analysis was performed using R software. For the beta-diversity analysis, Bray-Curtis dissimilarity between each pair of fecal microbiomes was computed and used for estimating the effect size of environmental factors using Adonis test. Statistical significance was determined: Student’s *t* test (comparison of two groups, normal distribution), one-way ANOVA (comparison of three or more groups). Wilcoxon rank-sum test or Wilcoxon signed-rank test was next used to determine the microbial features (such as microbial genera/species/strains and functional genes) that are enriched or depleted in a treatment group or a time point. The heatmap was built using the package “pheatmap” (v1.0.12). The R package “ggpubr” (v0.2.3) was used to make box plots. Spearman rank correlation was used to construct the co-occurrence network of molecular-level features (microbial taxa, functional genes and metabolites) features and clinical co-variates. Sankey diagram was further used to visualize the interplays among them using R package “ggalluvial”. The R package “gggenes” was used for visualization of the structure of *abfA* and *hypBA* gene clusters

## ACKNOWLEDGMENTS

This work was supported by the National Natural Science Foundation of China [No. 32021005, 32122067 and 31820103010].

## AUTHOR CONTRIBUTIONS

Principal investigators, C.C.Z, S.H, J.C.Z, and Q.X.Z.; metagenomic data analysis, C.C.Z, L.L.Y., C.C.M., and S.M.J.; metabolomic data analysis, C.C.Z., L.L.Y., and C.C.M.; bacterial isolation and genome sequencing and annotation: C.C.Z., and S.H.W.; phylogenetic and functional genomic analyses: C.C.Z., C.C.M., and L.L.Y; study design: F.W.T., J.X.Z., H.Z., W.C., L.M.L., Q.X.Z. and J.C.Z ; collection of samples and health information: Y.Z.X and C.C.Z.; manuscript drafting: C.C.Z., L.L.Y., Q.X.Z, S.H., and J.C.Z. All authors discussed the results, critically reviewed the text, and approved the final manuscript.

---

**Figure S1.**
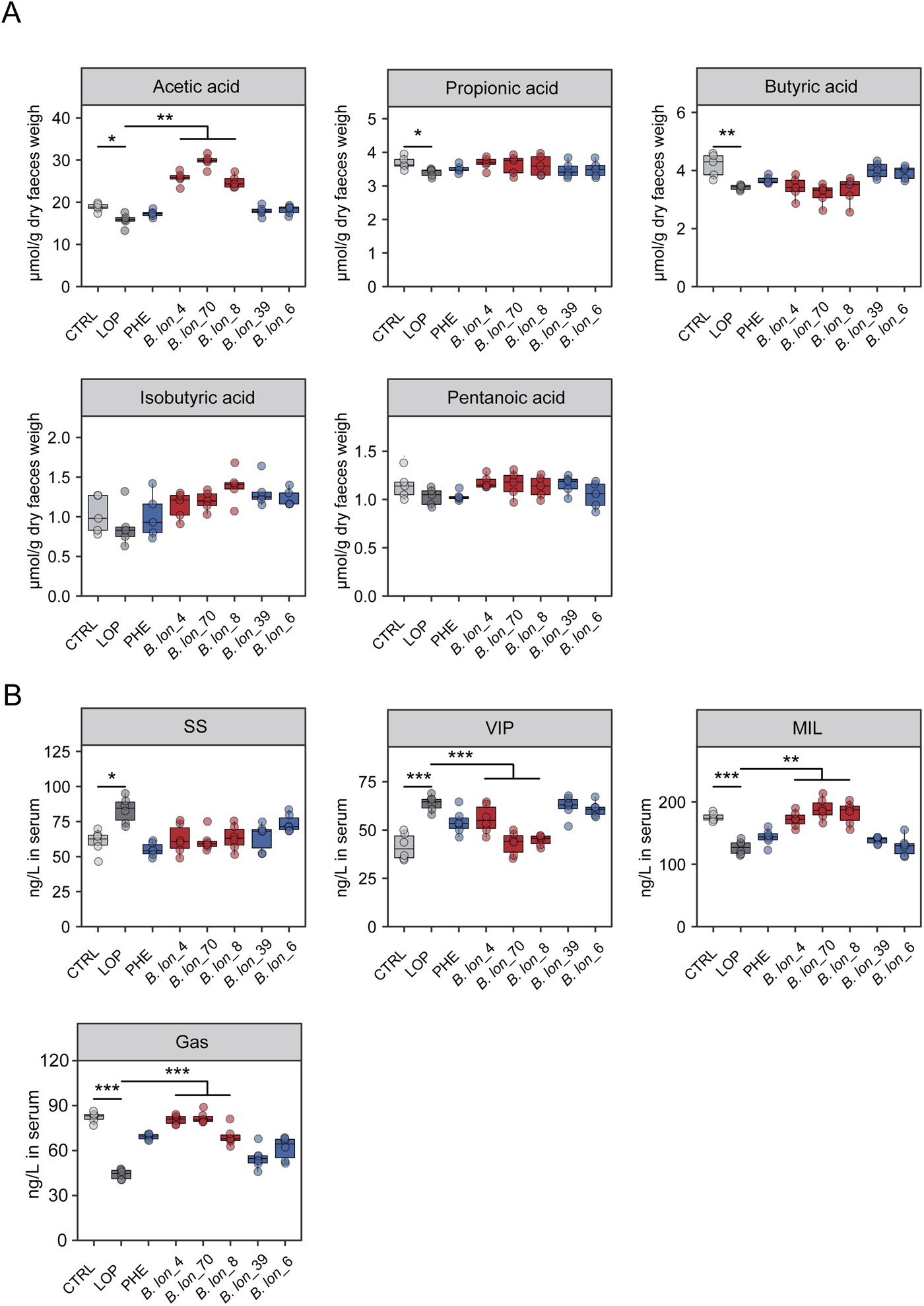
The abundance of short chain fatty acids and gastrointestinal peptide hormones in SPF mice determined with GC/MS and ELISA, respectively. (**A**) Concentrations of short chain fatty acids concentrations in mouse faces (N=4-6 biologically independent animals per group) determined with GC/MS. (**B**) MTL, Gas, SS and VIP levels in mouse serum determined with an enzyme-linked immunosorbent assay (ELISA). MTL, Motilin. Gas, gastrin. SS, somatostatin. VIP, vasoactive intestinal peptide. *p<0.05, **p<0.01, ***p<0.001 as determined by one-way ANOVA.

**Figure S2.**
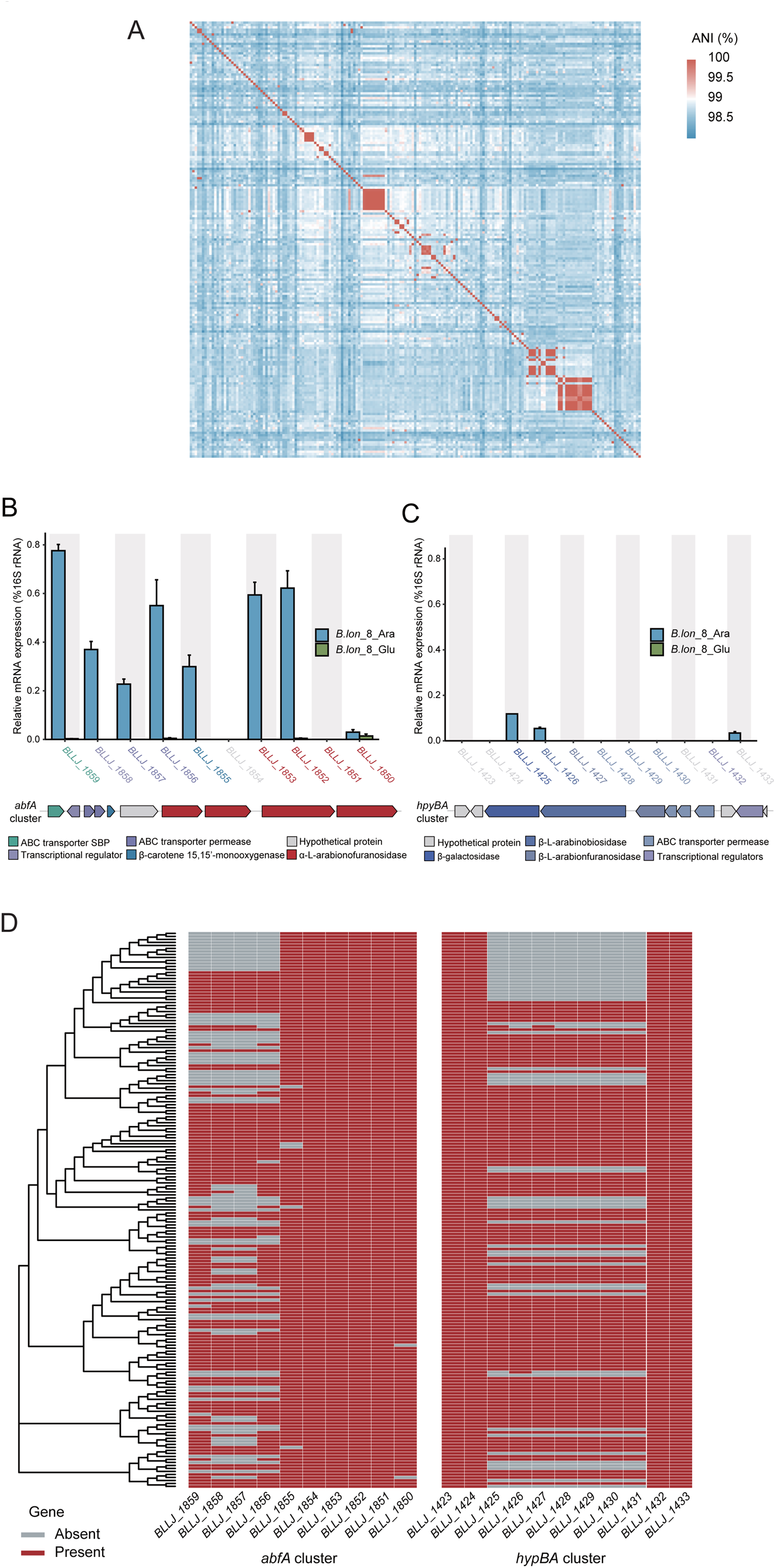
The genomic and transcriptomic features of effective and non-effective *B. longum* strains. Comparative genome analysis revealed that *B. longum* strains are similar to each other in genome-wide level and the distribution of the *abfA* and *hypBA* gene cluster are obviously different. Quantitative RT-PCR analysis revealed that the two gene clusters showed differently expressed induced by arabinan in *B. lon_8*. (**A**) A heatmap illustrates the ANI between each pair of 185 sequenced *B. longum* strains. (**B-C**) Expression level of functional genes in *abfA* cluster (**B**) and *hpyBA* cluster (**C**) of *B. lon_8* strain induced by arabinan *in vitro*. Bifidobacterial RNA was extracted from the *in vitro* culture after 12-hr incubation with addition of arabinan (Ara) or glucose (Glu) as a substrate. Gene expression levels are normalized as values relative to 16S rRNA expression. Error bars indicate SEM (N=3). (**D**) The presence or absence of *abfA* and *hypBA* gene clusters in each of 185 *B. longum* strains.

**Figure S3.**
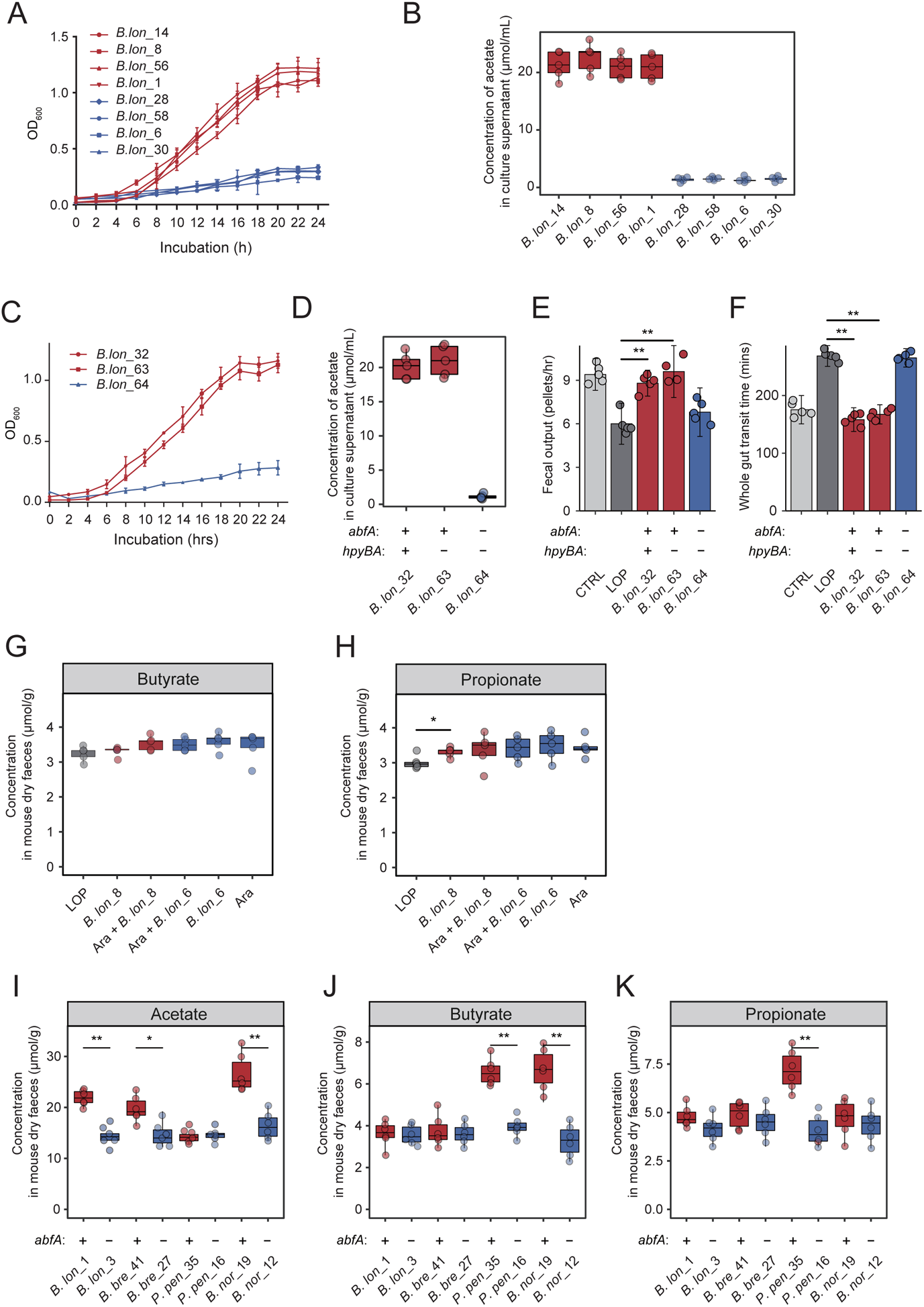
The impact of presence of *abfA* cluster in either *B. longum* strains on non-*B. longum* strains on the microbial phenotypes and metabolic profiles under *in vitro* or *in vivo* condition and corresponding host phenotypes (constipation index). (**A**) Growth of *B. longum* strains (*B. lon_*14, *B. lon_*8, *B. lon_*56, *B. lon_*1, *B. lon_*28, *B. lon_*58, *B. lon_*6, *B. lon_*1 and *B. lon_*30) in media containing arabinan. The strain was cultured at 37°C in sugar-restricted basal medium supplemented with 2.0% arabinan under anaerobic conditions. The OD_600_ was measured every 2 hrs for 24 hrs. Error bars indicate SEM (N=3). (**B**) Acetate concentrations in culture supernatant after 24 hrs incubation with addition of arabinan as a substrate, as determined by GC/MS. (**C**) Growth of *B. longum* strains (*B. lon_*32, *B. lon_*63 and *B. lon_*64) in media containing arabinan. The strain was cultured at 37°C in sugar-restricted basal medium supplemented with 2.0% arabinan under anaerobic conditions. OD_600_ was measured every 2 hrs for 24 hrs. Error bars indicate SEM (N=3). (**D**) Concentration of acetate in culture supernatant after 24 hrs incubation with addition of arabinan as a substrate, as determined by GC/MS. (**E-F**) Constipation related index from mice treated with *B. longum* harbors *abfA* gene cluster, *hypBA* gene cluster, or both. (**G-H**) The effects of *B. longum* strains and arabinan on concentrations of butyrate (**G**) and propionate (**H**) in constipation mouse model induced by loperamide. (**I-K**) The effects of bacteria strains on concentrations of acetate (**I**), butyrate (**J**) and propionate (**K**) in constipation mouse model induced by loperamide.

**Figure S4.**
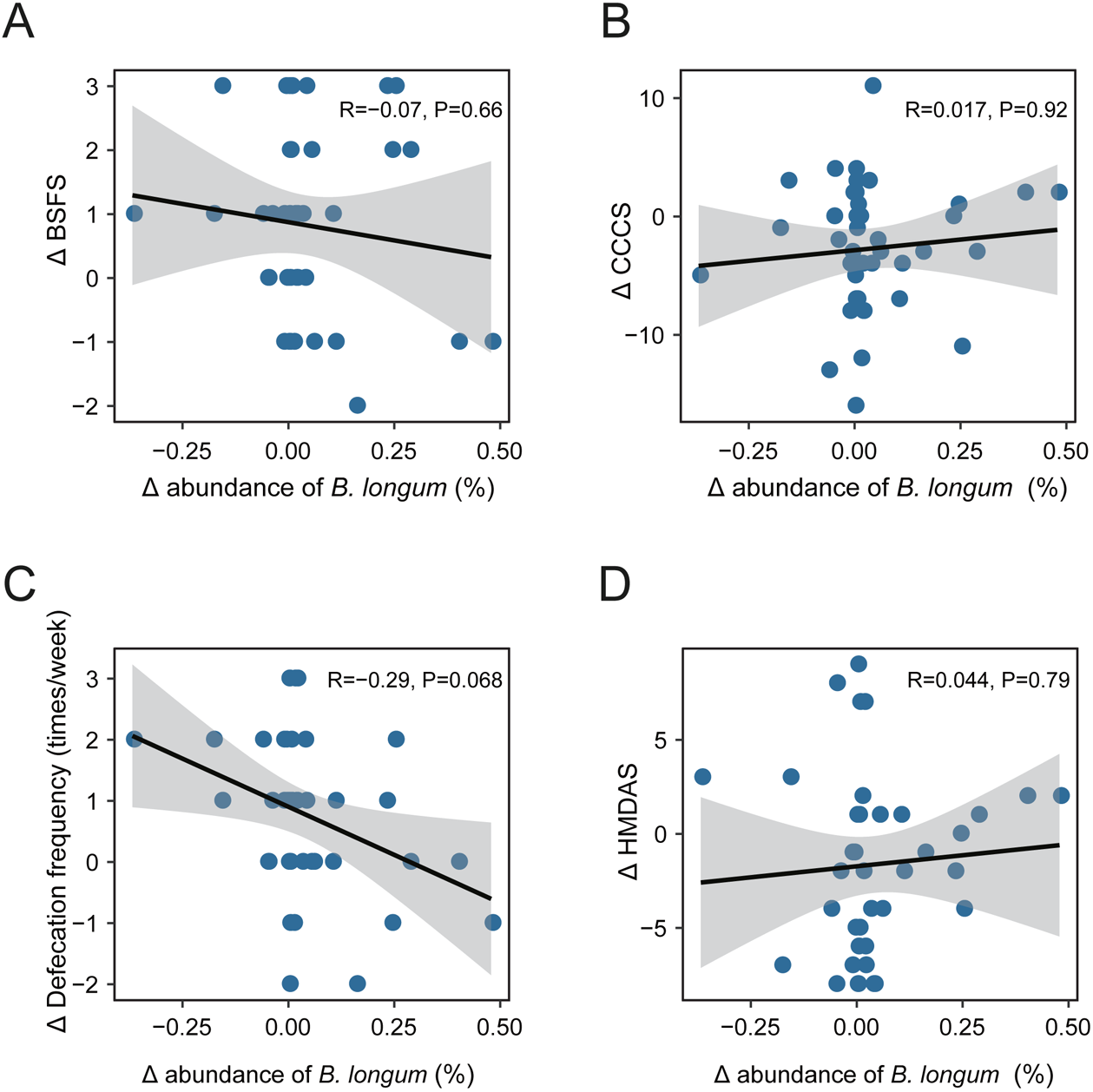
The change in clinical index has no association with the change in the overall abundance of fecal *B. longum* strains. (**A**) Spearman correlation between the change of BSFS and the change of *B. longum* abundance from hospital discharge to study completion. (**B**) Spearman correlation between the change of CCCS and the change of *B. longum* abundance from hospital discharge to study completion. (**C**) Spearman correlation between the change of defecation frequency and the change of *B. longum* abundance from hospital discharge to study completion. (**D**) Spearman correlation between the change of HMDAS and the change of *B. longum* abundance from hospital discharge to study completion. Spearman’s coefficients and p value for each correlation analysis are shown in (**A-D**).

**Figure S5.**
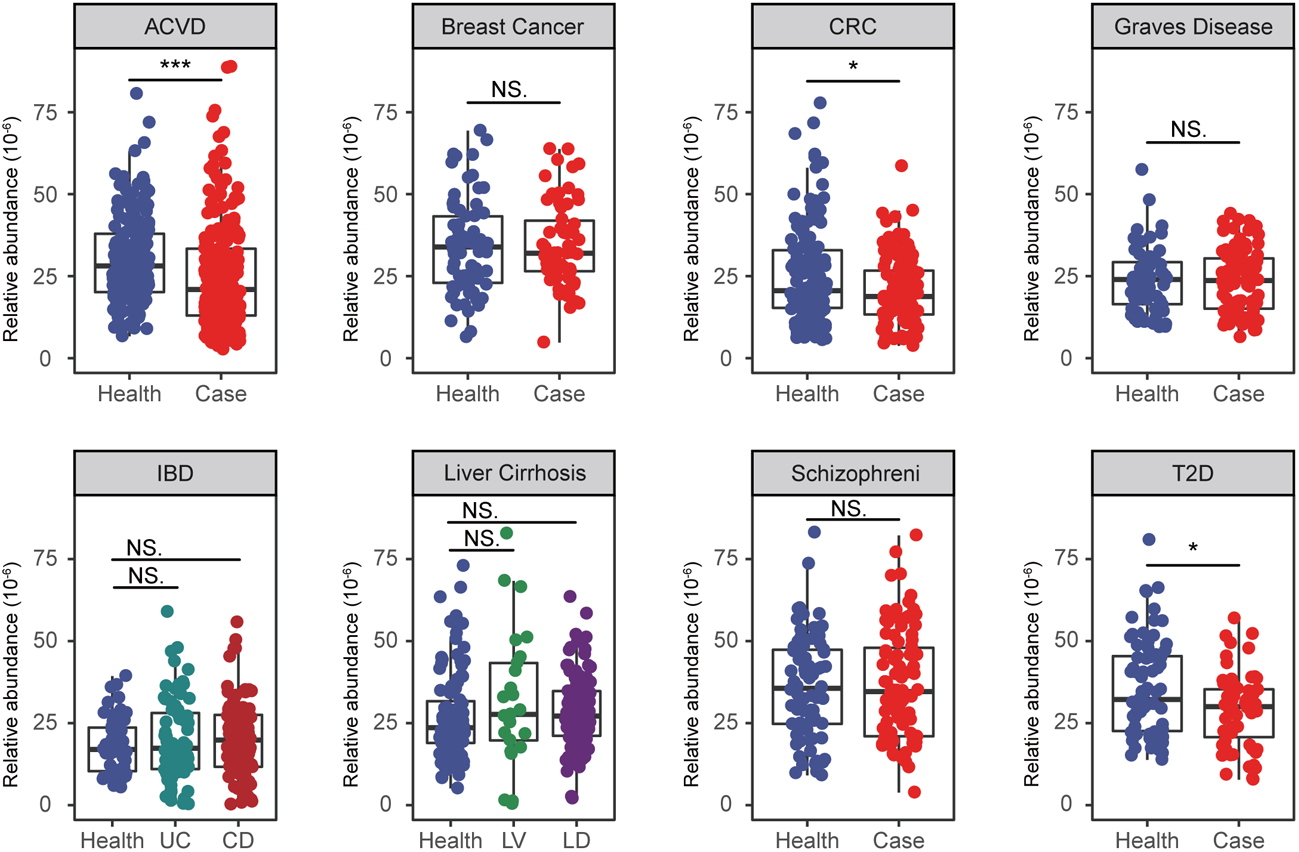
The abundance of *abfA* gene cluster can distinguish many other human disease phenotypes. The boxplots indicate the abundance of *abfA* gene cluster in case and control groups in each metagenomic dataset. ACVD: atherosclerotic cardiovascular disease, SCZ: Schizophrenia, GD: Graves’ disease, T2D: Type 2 diabetes, CRC: Colorectal cancer, BC: Breast cancer, UC: Ulcerative colitis, CD: Crohn’s disease, LC: Liver cirrhosis.

**Figure S6.**
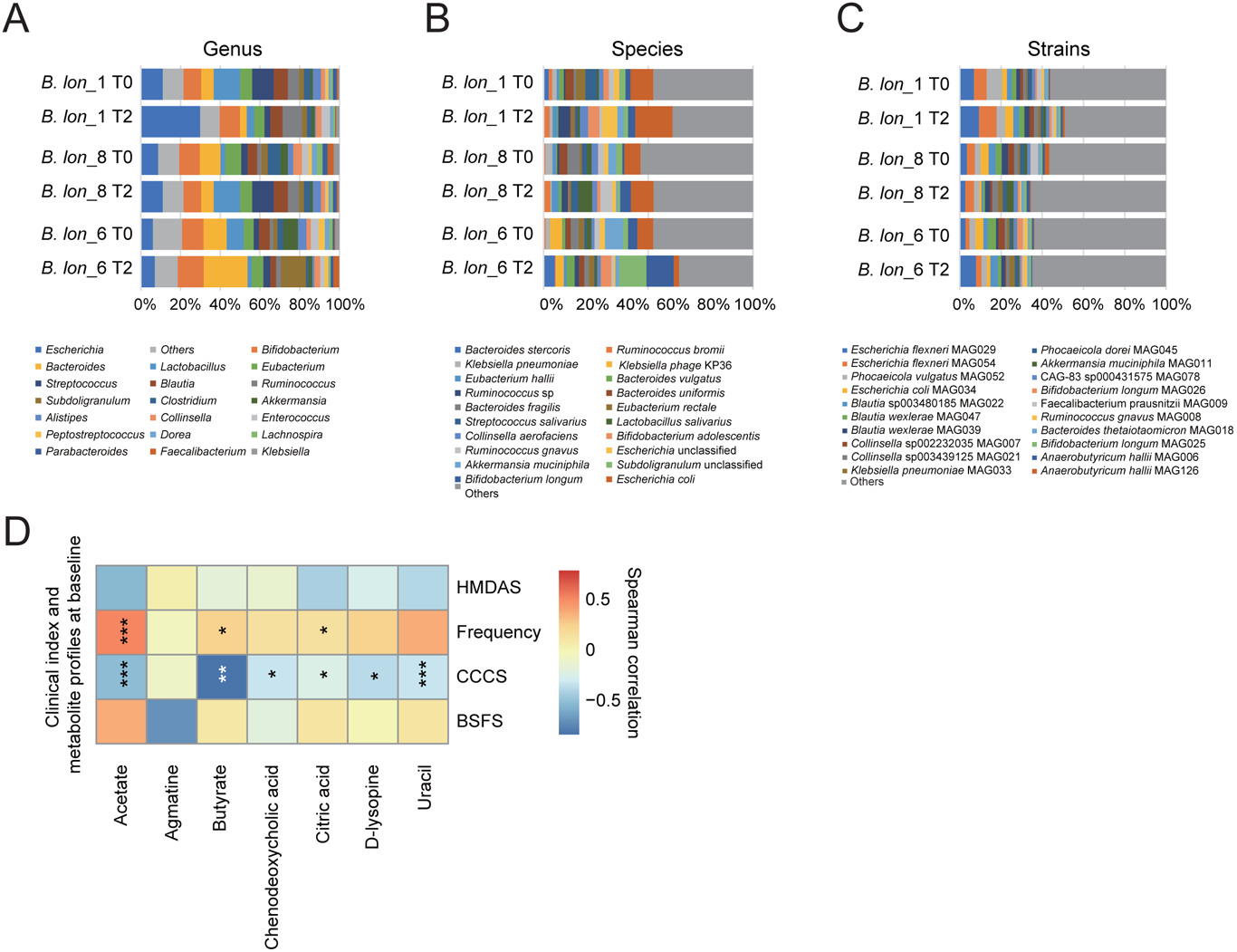
The alterations in fecal microbial composition in the mouse feces induced by *B. longum* administration. (**A-C**) Relative abundance of fecal microbial taxa was detected at the genus (**A**), species (**B**) and strains (**C**) level, for which only top-20 taxa in abundance are shown. (**D**) The CCCS inversely correlated with the relative abundance of selected fecal metabolites (acetate, butyrate, chenodeoxycholic acid and uracil). Symbols indicate significance: *p<0.05, **p<0.01, ***p<0.001.

**Figure S7.**
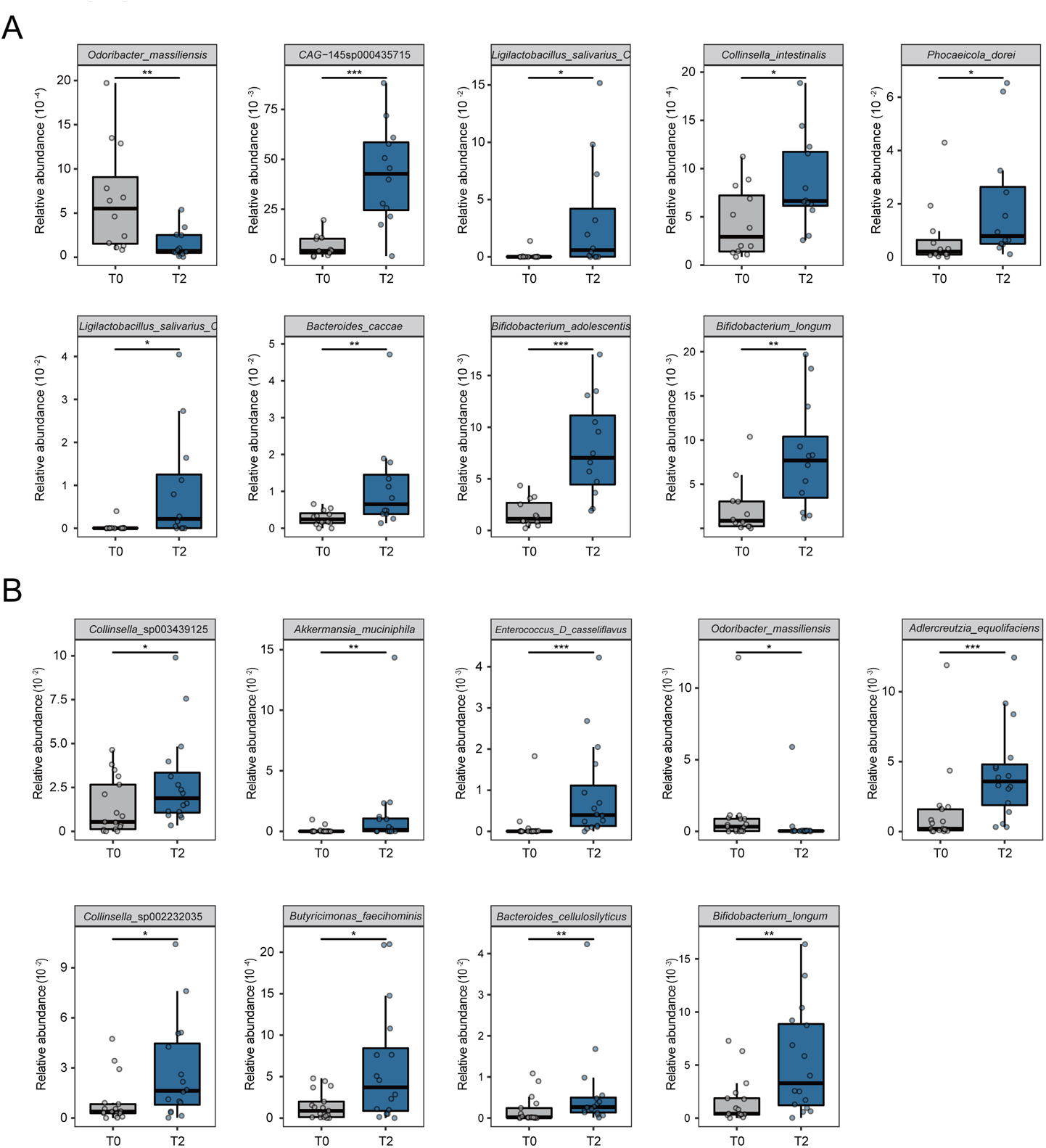
The differentially abundant microbial stains in the gut induced by the *B. lon_*1 and *B. lon_*8 administration. (**A**) The relative abundance of bacterial strains in mouse feces at baseline and 28 days after *B. lon_*1 administration. (**B**) The relative abundance of bacterial strains in mouse feces at baseline and 28 days after *B. lon_*8 administration. Only individuals with paired data were included in each analysis. Wilcoxon signed-rank tests were performed to detect the differential abundant microbes in *B. lon_*1 or *B. lon_*8 group before and after 28 days of supplementation. In the box plots, the line in the middle of the box corresponds to the median, and the inferior and superior limits of the box correspond to the 25th and the 75th percentiles, respectively. Symbols indicate significance: *p<0.05, **p<0.01, ***p<0.001.

